# *CNTN5*^−/+^ or *EHMT2*^−/+^ iPSC-Derived Neurons from Individuals with Autism Develop Hyperactive Neuronal Networks

**DOI:** 10.1101/368928

**Authors:** Eric Deneault, Muhammad Faheem, Sean H. White, Deivid C. Rodrigues, Song Sun, Wei Wei, Alina Piekna, Tadeo Thompson, Jennifer L. Howe, Leon Chalil, Vickie Kwan, Susan Walker, Peter Pasceri, Frederick P. Roth, Ryan K.C. Yuen, Karun K. Singh, James Ellis, Stephen W. Scherer

## Abstract

Induced pluripotent stem cell (iPSC)-derived cortical neurons are increasingly used as a model to study developmental aspects of Autism Spectrum Disorder (ASD), which is clinically and genetically heterogeneous. To study the complex relationship of rare (penetrant) variant(s) and common (weaker) polygenic risk variant(s) to ASD, “isogenic” iPSC-derived neurons from probands and family-based controls, for modeling, is critical. We developed a standardized set of procedures, designed to control for heterogeneity in reprogramming and differentiation, and generated 53 different iPSC-derived glutamatergic neuronal lines from 25 participants from 12 unrelated families with ASD (14 ASD-affected individuals, 3 unaffected siblings, 8 unaffected parents). Heterozygous *de novo* (7 families; 16p11.2, *NRXN1*, *DLGAP2*, *CAPRIN1*, *VIP*, *ANOS1*, *THRA*) and rare-inherited (2 families; *CNTN5*, *AGBL4*) presumed-damaging variants were characterized in ASD risk genes/loci. In three additional families, functional candidates for ASD (*SET*), and combinations of putative etiologic variants (*GLI3/KIF21A* and *EHMT2/UBE2I* combinations in separate families), were modeled. We used a large-scale multi-electrode array (MEA) as our primary high-throughput phenotyping assay, followed by patch clamp recordings. Our most compelling new results revealed a consistent spontaneous network hyperactivity in neurons deficient for *CNTN5* or *EHMT2.* Our biobank of iPSC-derived neurons and accompanying genomic data are available to accelerate ASD research.

## Introduction

The past two decades of research has determined Autism Spectrum Disorders (ASD) to be clinically (Fernandez and Scherer, 2017, Jones and Lord, 2013, Mahdi et al., 2018) and genetically (De Rubeis et al., 2014, Gilman et al., 2011, Pinto et al., 2014, Tammimies et al., 2015, Yuen et al., 2017) heterogeneous. Phenotypically, the fifth edition of the Diagnostic and Statistical Manual of Mental Disorders (DSM-5) combines autistic disorder, Asperger disorder, childhood disintegrative disorder and pervasive developmental disorder not otherwise specified into the single grouping of ASD (DSM-V, 2013). There are also syndromic forms of ASD (Carter and Scherer, 2013), and now more than 100 other disorders carrying different names (Betancur, 2011), that in a proportion of subjects can also present the necessary symptoms for an ASD diagnosis.

From the perspective of genetics, heritability estimates and family studies definitely demonstrate genes to be involved (Ronald and Hoekstra, 2011). Single high-penetrance genes and copy number variation (CNV)-affected loci, have now been implicated as *bona fide* autism-susceptibility (or risk) genes, although none of them show specificity for ASD alone (Malhotra and Sebat, 2012). These genetic alterations are rare in the population (<1% population frequency), and in some individuals, combinations of rare genetic variants affecting different genes can be involved (Devlin and Scherer, 2012), including more complex structural alterations of chromosomes (Brandler et al., 2018, Marshall et al., 2008). Recent research studying common genetic variants indicates polygenic contributors may be involved, and these can also influence the clinical severity of rare penetrant variants in ASD risk genes (Weiner et al., 2017).

Nearly 1,000 putative ASD risk loci are catalogued, with ~100 already being used in the clinical diagnostic setting (Carter and Scherer, 2013, Winden et al., 2018). There are some genotype-phenotype associations emerging, including general trends considering medical complications and IQ (Bishop et al., 2017, Sanders et al., 2015, Tammimies et al., 2015), sibling variability depending on the ASD gene variant they carry (Yuen et al., 2015), and lower adaptive ability in those carrying variants compared to affected siblings without the same genetic change (Yuen et al., 2017). Many of the ASD risk genes identified are connected into gene networks including those involved in synaptic transmission, transcriptional regulation, and RNA processing functions (Bourgeron, 2015, De Rubeis et al., 2014, Geschwind and State, 2015, Pinto et al., 2014, Sahin and Sur, 2015, Yuen et al., 2017, Yuen et al., 2016), with the impacted genes being involved in all of prenatal, region-specific, or broader brain development (Uddin et al., 2014). A general unifying theme that is emerging from neurophysiologic studies is an increased ratio of excitation and inhibition in key neural systems that can be perturbed by variants in the ASD risk genes, or environmental variables affecting the same targets (Canitano and Pallagrosi, 2017).

The advent of the induced pluripotent stem cell (iPSC) technology (Takahashi et al., 2007, Yu et al., 2007), followed by cellular re-programming to forebrain glutamatergic neurons (Habela et al., 2016), allows accessible cellular models to be developed for the highly heterogeneous ASD (Beltrao-Braga and Muotri, 2017, Dolmetsch and Geschwind, 2011, Durak and Tsai, 2014, Karmacharya and Haggarty, 2016, Marchetto et al., 2017, Yoon et al., 2014, Zhang et al., 2013). Carrying the precise repertoire of rare and common genetic variants as the donor proband, iPSC-derived neurons represent the best genetic mimic of proband neurons for functional and mechanistic studies. Induced differentiation can be achieved with high efficiency and consistency using transient ectopic expression of the transcription factor NEUROG2 (Ho et al., 2016, Zhang et al., 2013), and this has been shown useful in diverse phenotyping projects (Deneault et al., 2018, Pak et al., 2015, Yi et al., 2016). Proband-specific iPSC-derived neuronal cells indeed provide a useful model to study disease pathology, and response to drugs, but throughput (both iPSC-derived neurons and phenotyping) is low, with costs still high. As a result, so far, only a few iPSC-derived neuronal lines are typically tested in a single study.

Here, we develop a resource of 53 different iPSC lines derived from 25 individuals with ASD carrying a wide-range of rare variants, and unaffected family members. We also used clustered regularly interspaced short palindromic repeats (CRISPR) editing (Jinek et al., 2012, Ran et al., 2013) to create six “isogenic” pairs of lines with or without mutation, to better assess mutational impacts. Upon differentiation into cortical excitatory neurons, we investigated synaptic and electrophysiological properties using the large-scale multi-electrode array (MEA) as well as more traditional patch-clamp recordings. Numerous interesting associations were observed between the genetic variants and the neuronal phenotypes analyzed. We share our general experiences and the bioresource with the community. We also highlight one of our more robust findings—an increased neuronal activity in glutamatergic neurons deficient in one copy of *CNTN5* or *EHMT2*—which could be responsible for ASD-related phenotypes.

## Results

### Selection and Collection of Tissue Samples for Reprogramming

Participants were enrolled in the Autism Speaks MSSNG whole-genome sequencing (WGS) project (Yuen et al., 2017). All ASD and related control-participants were initially consented for WGS and upon return of genetic results, then consented for the iPSC study, using approved protocols through the Research Ethics Board at the Hospital for Sick Children. Some families were also examined by whole exome sequencing. The study took place over a 5-year period and used incrementally developing ASD gene lists from the following papers (Jiang et al., 2013, Marshall et al., 2008, Tammimies et al., 2015, Yuen et al., 2015) (Table 1). These primarily considered data from the MSSNG project, the Autism Sequencing Consortium (De Rubeis et al., 2014), and the Simons Foundation Autism Research Initiative (SFARI) gene list (discussion below). A diversity of different ASD-risk variants was targeted ranging in size from 1 nucleotide single nucleotide variants (SNV) to an 823 kb CNV (Figure 1 and Table 1; corresponding genomic coordinates in **Table S1**). Typically, one proband and one sex-matched unaffected member (control) per family were included (Figure 1). In total, 14 probands and 11 controls participated, of which 21 were males and 4 were females (Figure 1 and Table 1). Cells from either skin fibroblasts or CD34+ blood cells were collected for reprogramming into iPSCs (Figure 2A and Table 1).

**Table 1.**
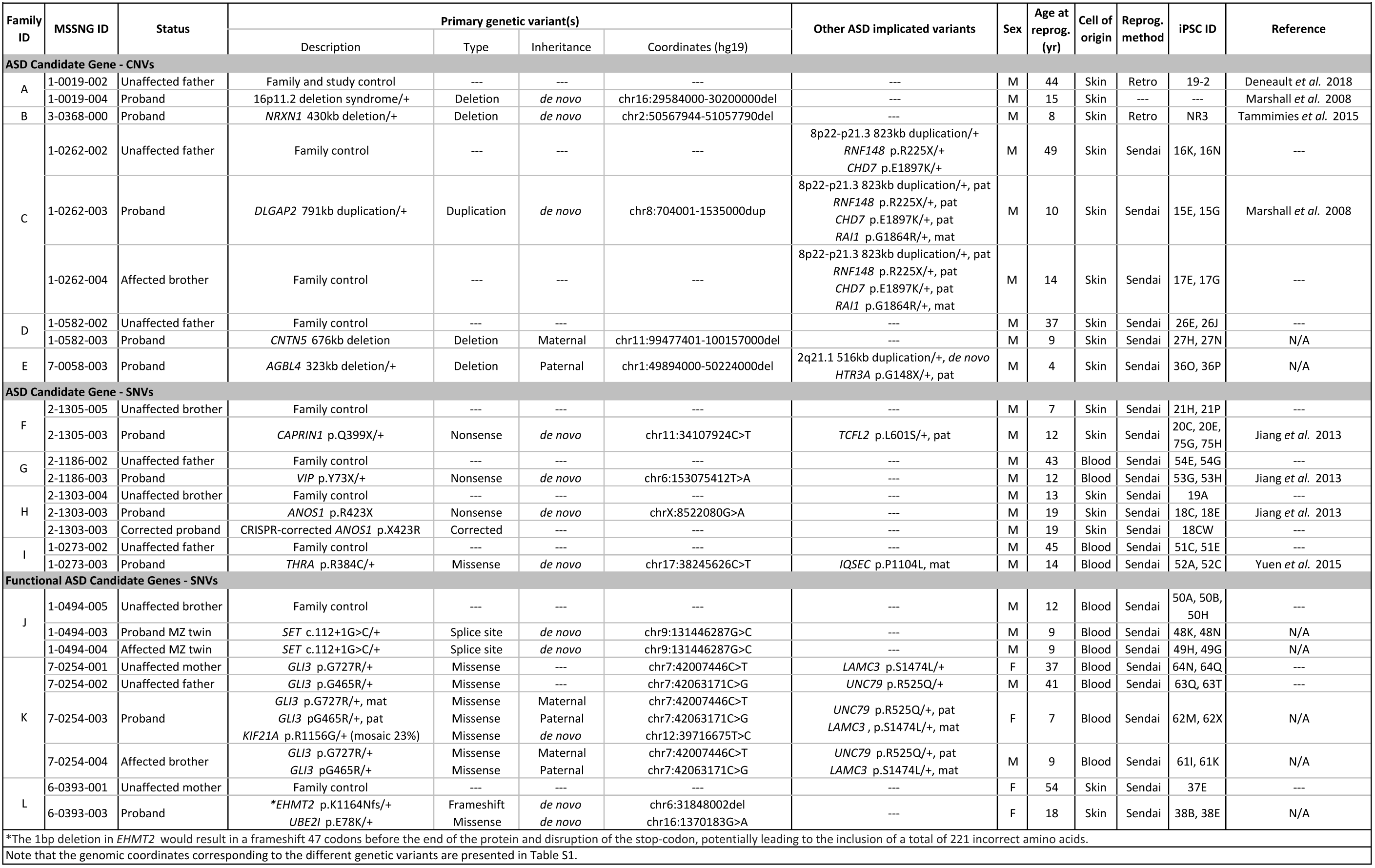
List of participants with ASD or unaffected controls, with the genetic variant(s) involved, and the different iPSC lines derived.

**Figure 1.**
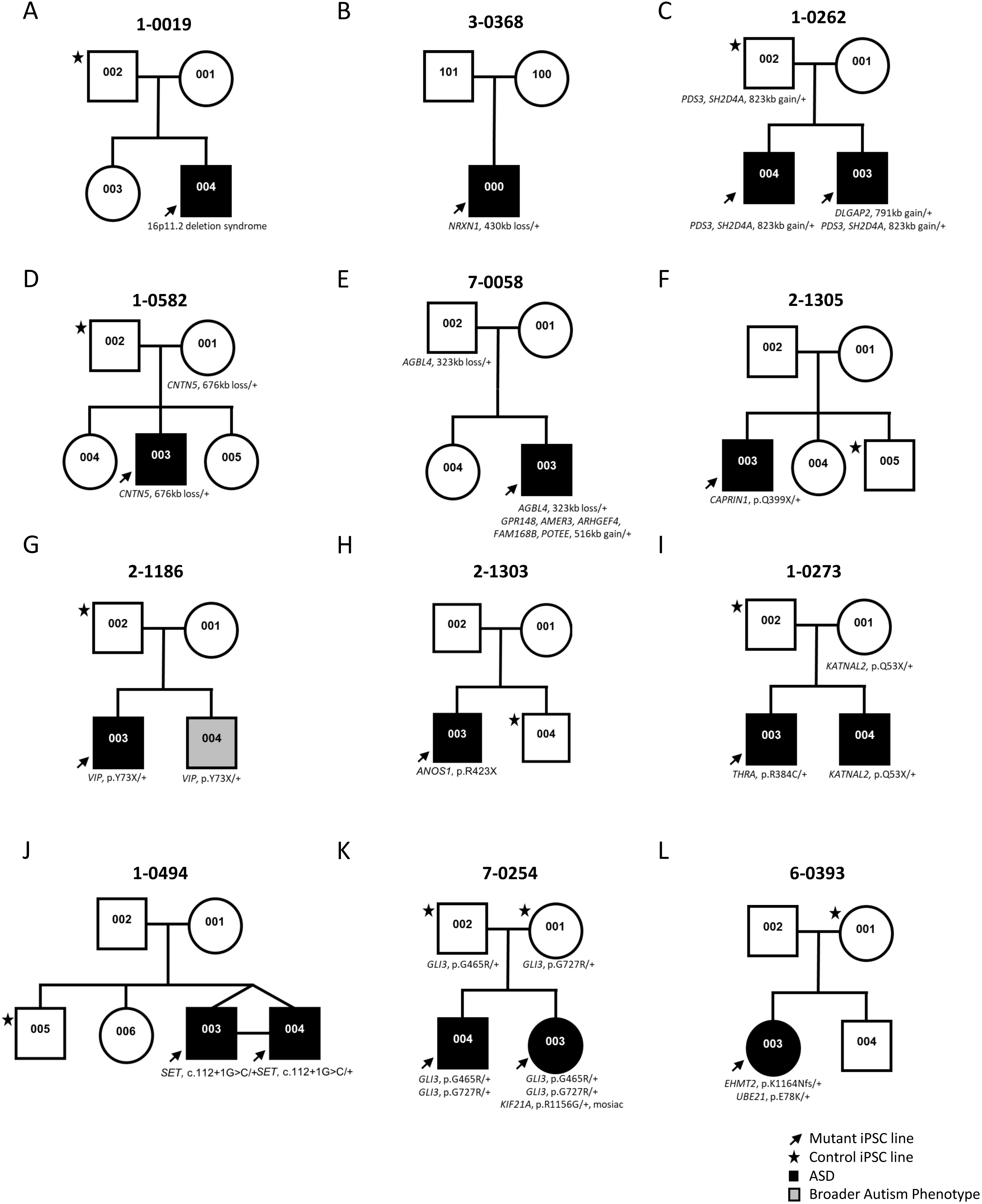
Genetic pedigrees of the participant families with identified genetic variants. One proband (black arrow) and one gender-matched unaffected member (black star) were typically selected for iPSC reprogramming. ASD-affected children are represented with a black box; note that line 1-0019-002 (19-2) in A) was used as a control and was described previously (Deneault et al., 2018).

**Figure 2.**
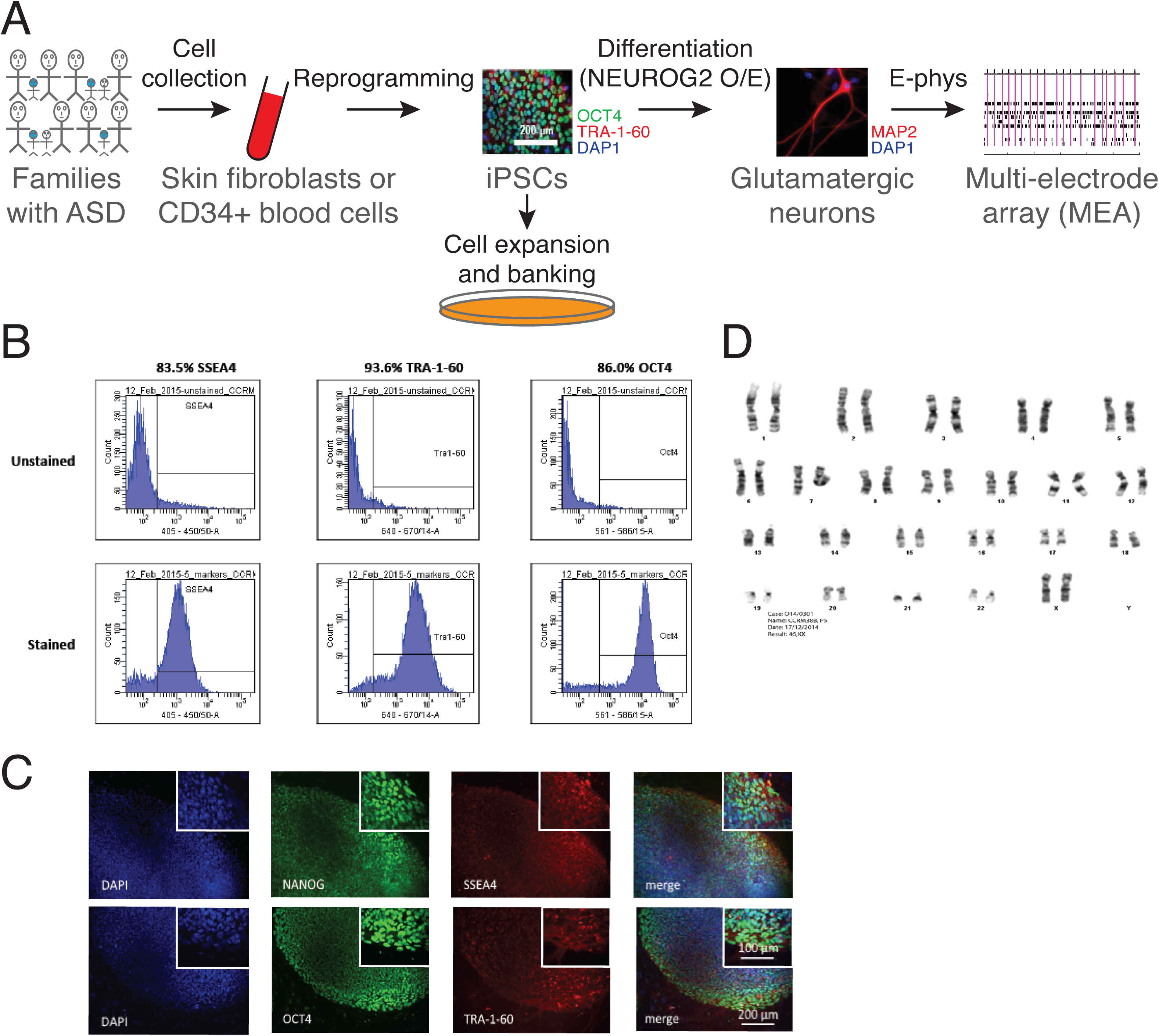
Generation of iPSCs and neurons. (A) Schematic representation of the experimental procedure to find specific electrophysiological signatures associated with genetic variants of clinical significance to autism spectrum disorder (ASD). Fibroblasts or blood cells were reprogrammed into iPSCs from a cohort of 25 probands and unaffected family members. Differentiation of iPSCs into glutamatergic neurons was achieved with NEUROG2 7-day transient overexpression, and electrophysiological properties were monitored using a multi-electrode array (MEA) device. (B) Flow cytometry and (C) Immunohistochemistry revealing expression of the pluripotency markers NANOG, SSEA4, OCT4 and TRA-1-60 in a representative iPSC line. (D) Representative normal male karyotype in iPSC; 20 cells were examined.

### Derivation of iPSC Lines

Two different viral approaches were used for cell reprogramming. For historical reasons, the first two lines in Table 1, namely 19-2 and NR3, were reprogrammed using retroviruses expressing *OCT4*/*POU5F1*, *SOX2*, *KLF4* and *MYC*, and a lentiviral vector that encoded the pluripotency reporter EOS-GFP/Puro^R^ (Hotta et al., 2009). Then, we moved to non-integrative Sendai virus for all the other tested lines (Table 1). Emerging iPSC colonies were selected for activated endogenous human pluripotency markers, differentiation potential into three germ layer cells after embryoid body formation *in vitro*, and normal karyotype (Figure 2B-D and **Table S2**). Two separate pluripotent and karyotypically normal iPSC lines were typically selected per participant for neuronal differentiation and phenotyping experiments (Table 1).

### Transient Induction of Neuronal Differentiation

We induced differentiation of newly generated iPSCs into glutamatergic neurons to test their electrophysiological properties (Figure 2A). We used the NEUROG2 ectopic expression approach since highly-enriched populations of glutamatergic neurons can be obtained within a week, and they exhibit robust synaptic activity when co-cultured with glial cells (Zhang et al., 2013). Importantly, we determined that this strategy offers highly uniform differentiation levels between cell lines derived from different participants (Deneault et al., 2018). This consistency was necessary to perform suitable phenotyping assays such as network electrophysiology recordings of several different lines in the same experimental batch. The resulting glutamatergic neurons were all subjected to electrophysiological phenotyping.

### Multi-Electrode Array Analysis of iPSC-Derived Neurons

MEA phenotyping was predominantly used in order to monitor the excitability of several independent cultured neuron populations in parallel, and in an unbiased manner. We sought to determine if any selected ASD-risk variants would interfere with spontaneous spiking and synchronized bursting activity in a whole network of interconnected glutamatergic neurons. We ensured that the duration and amplitude of detected spikes were similar to typical mammalian neurons, i.e., action potential widths of around 1-2 milliseconds (ms) and peak amplitudes of approximately 20-150 μV (**Figure S1A**). We monitored the weighted mean firing rate (MFR), which represents the MFR per active electrode, four to eight weeks post-NEUROG2 induction (PNI), in all control cells, i.e., all unaffected sex-matched family members from all tested families. After pooling all control samples together, the highest MFR was observed at week six PNI (**Figure S1A**, right panel). Hence, we opted to use week six PNI as the common reading time point for comparison of all cell lines.

We measured the glutamatergic/GABAergic nature of our cultured neurons produced using NEUROG2 ectopic expression, which is known to repress GABAergic differentiation at the advantage of glutamatergic (Roybon et al., 2010). MFRs were measured upon treatment with different receptor inhibitors. No substantial change was observed after addition of the GABA receptor inhibitor PTX (**Figure S1B**), indicating that GABAergic neurons are not appreciably present in our cultures. However, the MFR was significantly reduced in the presence of the AMPA receptor inhibitor CNQX while unchanged in untreated cells (**Figure S1B**). This further suggests that most of the culture was composed of glutamatergic neurons. All activity was abolished after addition of the sodium channel blocker TTX (**Figure S1B**), indicating that our human neurons were expressing functional sodium channels.

Based on these control experiments, we examined our proband lines, starting with those identified with a simple set of putative ASD-related variants, e.g., *NRXN1, CNTN5*, *CAPRIN1*, *VIP, ANOS1, THRA* and *SET* (Figure 1 and Table 1). Two main MEA metrics were monitored, i.e., weighted MFR and network burst frequency, at week six PNI. The heterozygous *CNTN5*-mutant neuron lines 27H and 27N presented a significantly higher weighted MFR compared with their related *CNTN5^+/+^* control lines 26E and 26J (Figure 3A), indicating an increased spontaneous spiking activity in glutamatergic neurons. These two lines also showed a significantly higher network burst frequency (Figure 3A), indicating a more synchronized neuronal activity across each well. Importantly, CNTN5 protein levels were reduced by at least 33% in *CNTN5*^−/+^ neurons (Figure 3A), suggesting that the 676-kb heterozygous loss in *CNTN5* interferes with the expression of CNTN5 protein and may be responsible for higher levels of neuronal activity.

**Figure 3.**
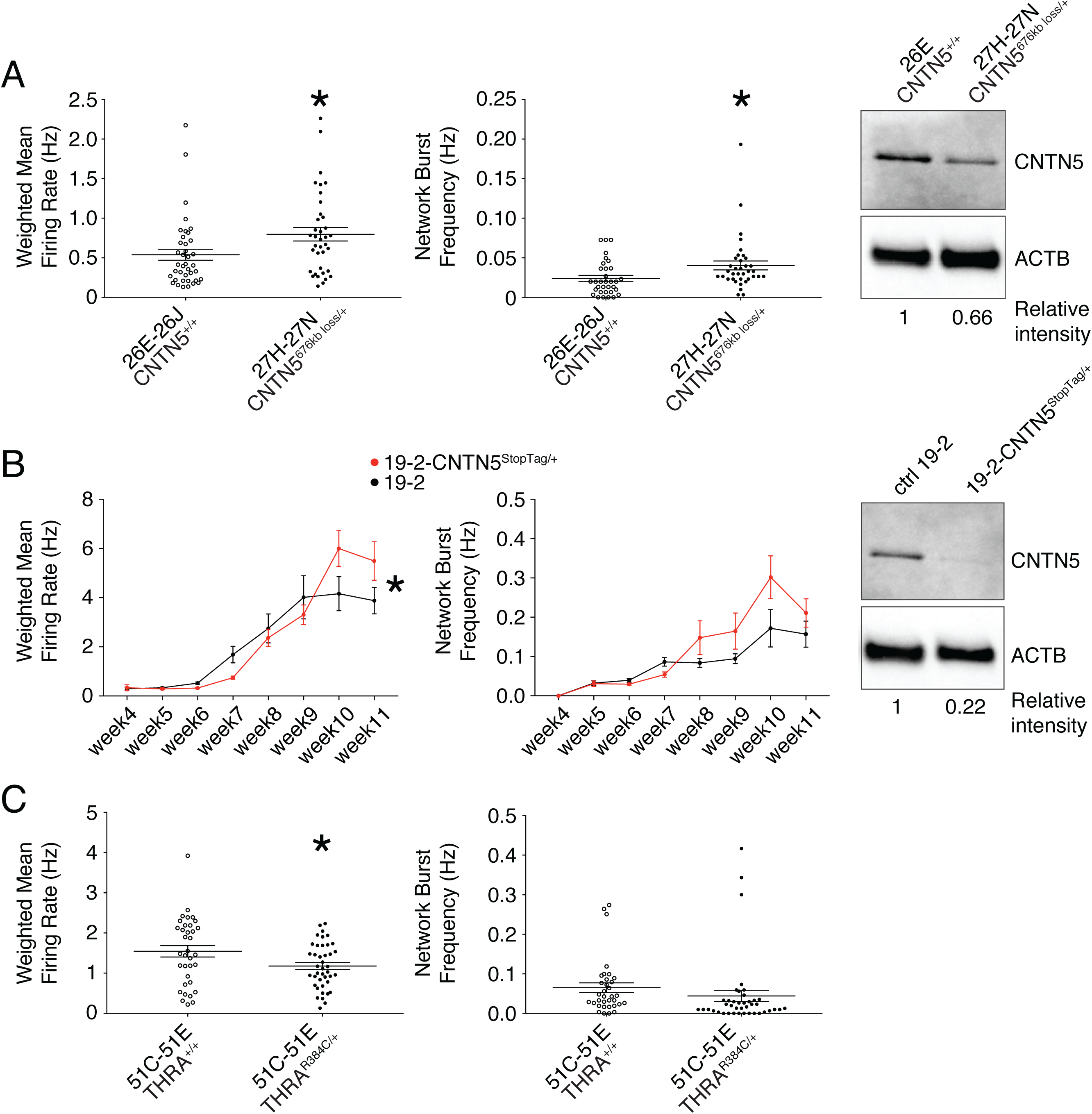
Multi-electrode array monitoring of iPSC-derived glutamatergic neurons. Weighted mean firing rate (MFR) and network burst frequency were recorded from (A) CNTN5 family at week 6 PNI, (B) CNTN5-isogenic pair 19-2 from week 4-11 PNI, and (C) THRA family NEUROG2-neurons at week 6 PNI; weighted MFR represents the MFR per active electrode; an electrode was considered active when a minimum of 5 spikes per minute were detected; a network burst was identified as a minimum of 10 spikes with a maximum inter-spike interval of 100 ms, occurring on at least 25% of electrodes per well; iPSC IDs and genotypes are indicated below each graph; values are presented as mean±SEM of two different lines per participant, and of several technical and biological replicates, as presented in Table S3. Right panels show western blots revealing protein levels in mutant and control iPSC-derived neurons; actin beta (ACTB) was used as a loading control and the relative intensity of each band is indicated below the blots. * p < 0.05 from unpaired t test two-tailed (A and C), and from multiple t test comparison (B)

### Isogenic Pair to Control for Genetic Background Contribution

Unaffected gender-matched family members are genetically similar to their related probands, but still present substantial genetic differences that can contribute to a given phenotype. Isogenic cell pairs represent better control of the genetic background contribution (Hoffman et al., 2018). CRISPR editing provides the possibility to engineer such isogenic controls (Miyaoka et al., 2014, Powell et al., 2017). Since editing large CNVs, such as the 676-kb deletion in *CNTN5*, is currently limited by existing technology, we elected to introduce a set of nonsense mutations, previously described as “StopTag” (Deneault et al., 2018), to knock out (KO) the expression of this gene in an unrelated iPSC line that was previously generated from a non-ASD and non-carrier individual. This parental line “19-2” was also exploited in similar isogenic KO approaches (Woodbury-Smith et al., 2017a) (Ross et al., submitted; Zaslavsky et al., in revision), allowing assessment in a different and unrelated genetic background. For technical reasons, we targeted exon 5 of the transcript ENST00000524871.5 of *CNTN5* in order to disrupt its expression. A heterozygous iPSC line was isolated to better mimic the heterozygous status of the *CNTN5* deletion in the proband lines 27H and 27N. Intriguingly, the new isogenic iPSC-derived neuron line 19-2-*CNTN5*^StopTag/+^ did not show a significant difference in terms of both weighted MFR and network burst frequency at week six PNI (Figure 3B), suggesting that this different genetic background might contribute to the result. For this validation set up, we decided to record for longer times, i.e., until 11 weeks PNI. Interestingly, line 19-2-*CNTN5*^StopTag/+^ presented a significant higher weighted MFR at 10 weeks PNI compared to its control 19-2 (Figure 3B). Moreover, CNTN5 protein levels were clearly decreased in this isogenic mutant line (Figure 3B, right panel), implying that StopTag insertion efficiently disrupted gene expression. These results indicate that loss of CNTN5 function is responsible for increased neuronal activity *in vitro*.

By contrast, iPSC-derived neurons carrying the missense variant R384C in THRA presented a significantly decreased weighted MFR (Figure 3C), suggesting that this variant affects the function of THRA protein and limits neuronal activity. No other simple-variant lines presented significant difference in MEA results compared to their related controls (**Figure S2**). No unaffected familial control was available to compare with the line NR3, carrying a 430-kb loss in *NRXN1* (Figure 1B and Table 1). For this reason, we averaged the results obtained from all unrelated controls used in this study, compared it with line NR3, and obtained no significant differences (**Figure S2A**). Although a proper genetically-matched control was not available for line NR3, this suggests that the loss of one *NRXN1* allele does not interfere with control neuronal activity in this setting.

### Repair of *ANOS1* Rescues Defective Membrane Currents

In a complementary approach to minimize the confounding effects of genetic background from familial and unrelated controls, and its impact on phenotype, we sought to edit our proband-specific variants using CRISPR in order to create matching isogenic controls. We prioritized genes affected by nonsense variants such as *CAPRIN1*, *VIP* and *ANOS1* (Figure 1F-H and Table 1). For instance, the nonsense variant R423X found in *ANOS1* in participant 2-1303-003 was successfully corrected in the corresponding iPSC line 18C (Figure 4A-C). Indeed, after detecting 7% edited cells using droplet digital PCR (ddPCR) in well G08 in the primary 96-well culture plate post-nucleofection, two subsequent limiting-dilution enrichment steps were necessary to isolate a 100% corrected iPSC line (Figure 4B). Sanger sequencing confirmed the properly corrected genomic DNA sequence (Figure 4C). This newly corrected line was named “18CW” (see iPSC line ID “18CW” in Table 1 and Figure 4C).

**Figure 4.**
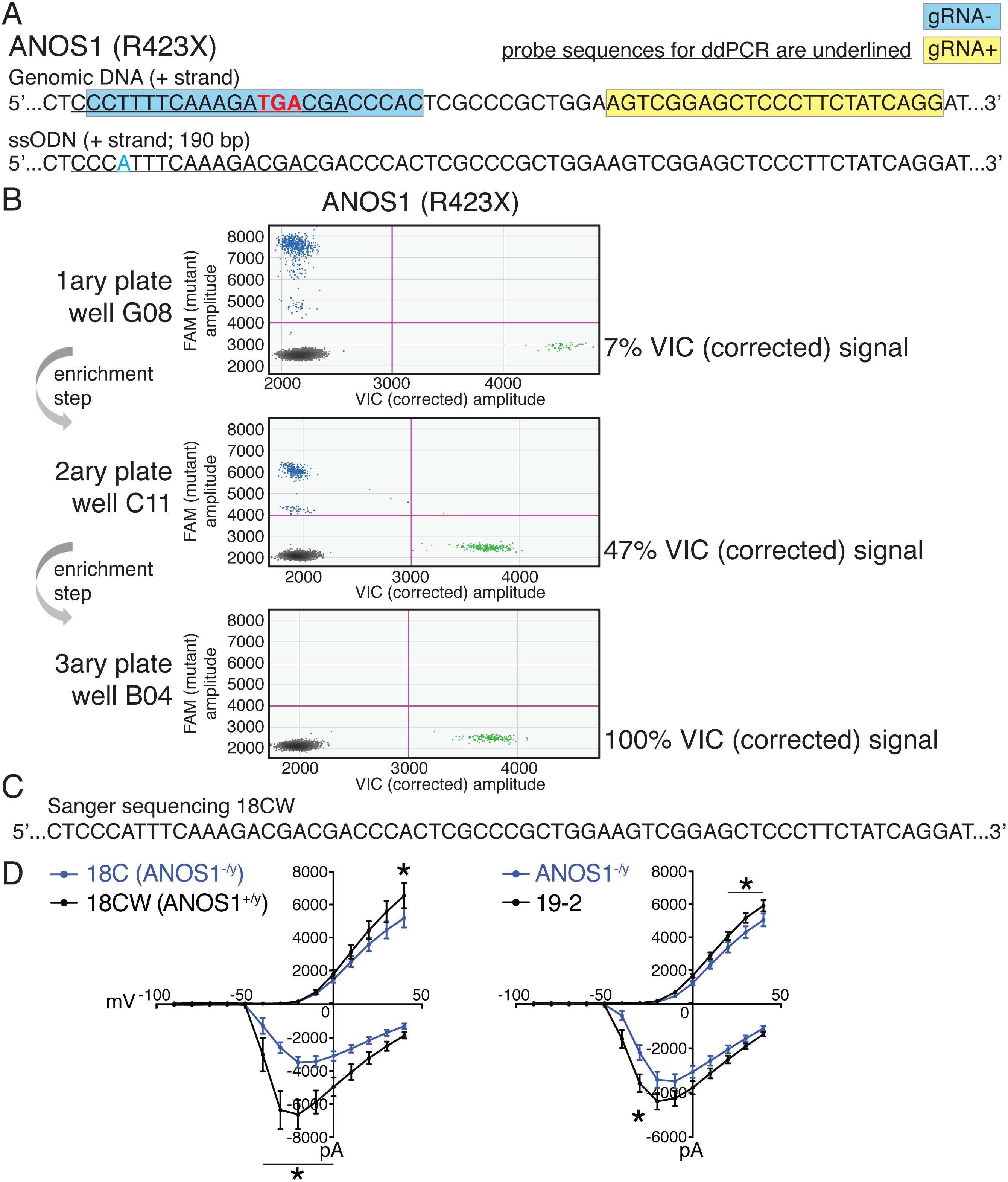
Correction of point mutation in ANOS1 in iPSCs using CRISPR editing. (A) Design of gRNAs, ssODNs and ddPCR probes for correction of R423X in ANOS1; one sgRNA for each genomic DNA strand, i.e., gRNA-in blue and gRNA+ in yellow, was devised in close proximity for the double-nicking system using Cas9D10A; the non-sense mutations in ANOS1 is depicted in bold red; a silent mutation was introduced in ssODN (in blue) for ddPCR probe (underlined) specificity and to prevent nicking. (B) ddPCR absolute quantification coupled with two consecutive limiting-dilution enrichment steps were necessary to isolate a 100% corrected line, i.e., 100% VIC signal. (C) Sanger sequencing confirmed proper correction of non-sense mutation R423X in line 18C back to wt; this newly corrected line was named 18CW. (D) Outward and inward membrane current detected by patch-clamp recordings; total number of recorded neurons was 15 for both 18C and 18CW, 20 for ANOS1^−/y^ and 33 for control 19-2; values are presented as mean±SEM of three independent differentiation experiments, recorded at day 21-25 PNI. * p < 0.05 from multiple t test comparison

Despite a behaviour similar to the unaffected familial control line 19A in terms of weighted MFR and network burst frequency, the CRISPR-corrected line 18CW did not exhibit a statistical difference from to its isogenic counterpart 18C (**Figure S2D**). Nonetheless, the availability of such isogenic set prompted us to explore more detailed electrophysiological properties using patch-clamp recordings of single neurons in order to reveal any phenotype not detected using MEA. While the advantage of MEA experiments is that continuous live monitoring of neural activity can be measured over multiple weeks, we used patch-clamp electrophysiology on NEUROG2 neurons between days 21-25 PNI. This time window provides robust recordings to detect phenotypes, as shown in previous studies (Yi et al., 2016). Furthermore, the increased density of neuronal processes appearing beyond 4 weeks PNI can preclude consistent clean patch-clamp recordings, but this is not an issue with MEA. Using this protocol, we detected significantly lower outward membrane current at 40 mV in the mutant line 18C compared to its isogenic control 18CW (Figure 4D, left panel). A significantly higher inward current was also observed in mutant neurons between-40 and 0 mV (Figure 4D, left panel). A comparable pattern of outward/inward current was monitored in a different unrelated isogenic pair (Figure 4D, right panel), in which our StopTag fragment was previously inserted within the *ANOS1* coding sequence in line 19-2 (Deneault et al., 2018). These results indicate that *ANOS1*-null iPSC-derived glutamatergic neurons present abnormal sodium and potassium membrane currents that might contribute to ASD development.

Notably, these observations underline that some specific electrophysiological phenotypes at the single cell level, e.g., membrane currents, may not be captured when using MEA monitoring at the cell population level.

### Addressing Complex-Variant Lines

Studying the relationship between ASD in individuals carrying multiple ASD-relevant variants requires additional considerations. For example, lines 36O and 36P were reprogrammed from participant 7-0058-003 and were each carrying a 323-kb deletion disrupting *AGBL4*, a 243-kb deletion disrupting *TCERG1L*, and a 516-kb duplication encompassing the *GPR148*, *AMER3*, *ARHGEF4*, *FAM168B*, *POTEE* genes (Figure 1E and Table 1). As for line NR3, no unaffected family member was available as control for lines 36O and 36P, thus we used the “all control average” and no significant differences were observed in terms of weighted MFR and network burst frequency (**Figure S3A**). No significant MEA recording differences were observed with *DLGAP2* (**Figure S3B**) or *GLI3/KIF21A* (**Figure S3C**) complex-variant mutant lines with respect to their corresponding familial controls. These results suggest that different expression levels of these genes do not interfere with control neuronal activity.

### Neuronal Hyperactivity in *EHMT2/UBE2I* Complex-Variant Neurons

Lines 38B and 38E from participant 6-0393-003 carry two ASD-relevant variants; a missense E78K in *UBE2I* and a frameshift variant K1164Nfs in *EHMT2* (Figure 1L and Table 1). MEA recordings between weeks 4-8 PNI showed a significantly higher weighted MFR and network burst frequency compared to their familial control line 37E (Figure 5A). To ensure that this hyperactivity was synaptic and not only intrinsic to the neurons, we performed patch-clamp recordings, at day 21-25 PNI to avoid the increased density of neuronal processes that impacts the ability to obtain clean recordings, as stated previously. Intrinsic properties, e.g., capacitance and resistance, did not vary significantly (Figure 5B), indicating comparable maturity levels between lines 37E and 38E. While spontaneous excitatory post-synaptic current (sEPSC) amplitude was unchanged, sEPSC frequency was significantly higher in mutant neurons compared to controls (Figure 5B). These observations suggest that a potential loss-of-function of *UBE2I* and/or *EHMT2* is involved in ASD-related neuronal dysfunction.

**Figure 5.**
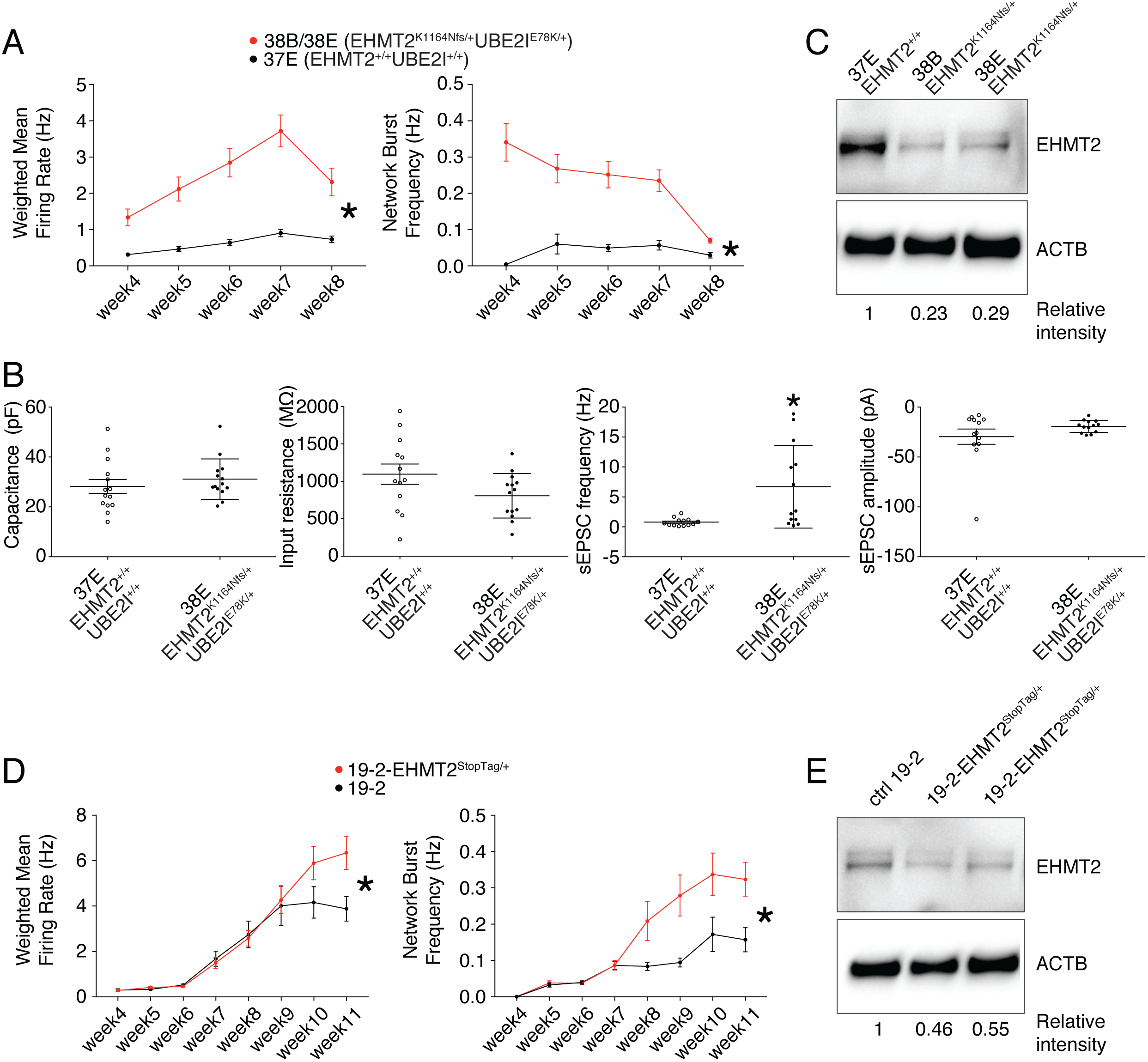
Electrophysiological and protein levels variations in EHMT2-deficient neurons. (A) Weighted mean firing rate (MFR) and network burst frequency were recorded using MEA from the EHMT2/UBE2I family from week 4-8 PNI; values are presented as mean±SEM of several technical and biological replicates, as presented in Table S3; * p < 0.05 from multiple t test comparison. (B) Patch-clamp recordings of two selected lines, i.e., 37E (control) and 38E (mutant); values are presented as mean±SEM of 14 different neurons from two independent differentiation experiments. * p < 0.05 from from unpaired t test two-tailed. (C) Western blot showing a decrease in EHMT2 protein levels in mutant neurons (38B and 38E) compared to their respective control neurons (37E); actin beta (ACTB) was used as a loading control and the relative intensity of each band is indicated below the blots. (D) MEA recordings of the isogenic pair 19-2 and 19-2-EHMT2^StopTag/+^ iPSC-derived neurons from week 4-11 PNI; values are presented as mean±SEM of eight different wells for each three independent differentiation experiments; * p < 0.05 from multiple t test comparison. (E) Western blot showing a decrease in EHMT2 protein levels in mutant neurons 19-2-EHMT2^Stop-Tag/+^ compared to their respective control (ctrl) neurons 19-2; ACTB was used as a loading control and the relative intensity of each band is indicated below the blots. pF = picofarad; MΩ = megaohm; Hz = hertz; pA = picoampere

### Evidence of Functional Impact of *EHMT2*, but not *UBE2l* Variants

Since our attempts to edit the variants E78K in *UBE2I* and K1164Nfs in *EHMT2* had not been successful, we sought to determine the potential contribution of E78K in *UBE2I* to the observed synaptic hyperactivity. To estimate the damaging potential of this missense variant on the function of UBE2I protein, we utilized a *Saccharomyces cerevisiae* complementation assay that was previously developed as a validated surrogate genetic system to predict the pathogenicity of diverse human variants (Sun et al., 2016). In this assay, lethality of a temperature-sensitive allele of the yeast *UBC9* gene (ortholog of human *UBE2I*) is rescued by expressing a functional version of human UBE2I. Several missense variants in *UBE2I* have been accurately predicted as deleterious at conserved positions, or benign at other positions (Zhang et al., 2017). Therefore, we used this complementation assay to test the consequence of our variant E78K, and found no effect of this variant on the function of human UBE2I (**Figure S4**). Because these results disfavor involvement of the *UBE2I* variant E78K in the neuronal hyperactivity observed in Figure 5A-B, we excluded *UBE2I* from subsequent experiments and further explored a potential causal link between *EHMT2* and synaptic activity.

Interestingly, evaluation of EHMT2 protein abundance revealed a clear decrease in the mutant lines 38B and 38E, as compared to the control 37E (Figure 5C). This suggests that a reduced expression of *EHMT2* increases spontaneous spiking activity and sEPSC frequency of glutamatergic neurons.

### *EHMT2*^−/+^ CRISPR-Isogenic Pair Confirms Neuronal Hyperactivity

Since the prediction of damage extent of the frameshift variant K1164Nfs on the function of EHMT2 may not be accurate, we used our StopTag insertion strategy in iPSC line 19-2, and targeted exon 20 of the transcript ENST00000375537.8 of *EHMT2* in order to disrupt its expression. In this new isogenic line, weighted MFR and network burst frequency were also significantly increased in iPSC-derived 19-2-*EHMT2^StopTag^*^/+^ neurons compared to control 19-2, around week 9 PNI and beyond (Figure 5D). This increased activity in mutant neurons occurred later than that observed in the familial lines 38B/E, possibly due to different genetic backgrounds. Accordingly, EHMT2 protein levels were reduced by half in mutant cells (Figure 5E). We also performed patch-clamp recordings on these neurons at day 21-25 PNI, as above. We did not detect any significant change in sEPSC frequency and amplitude at this earlier time point compared to the MEA experiment. However, intrinsic properties showed a significant increase in capacitance and a lower resting membrane potential in mutant cells (**Figure S5**). These observations suggest that the mutant neurons at 3-4 weeks PNI potentially have a faster maturation rate, however this phenotype is most pronounced in the hyperactivity recorded by MEAs later at 9-11 weeks PNI. These results support the conclusion that inactivation of one allele of *EHMT2* significantly increases spontaneous network activity of excitatory neurons, with possible effects on the neuronal maturation process.

## Discussion

In order to establish a scalable iPSC-derived neuron paradigm to study ASD, we selected 12 well-characterized families bearing assumed etiologic variants in ASD-relevant genes, and loci. Per family, we established one to four different fully-characterized and normal iPSC lines from typically one individual with ASD, and one unaffected (non-ASD) gender-matched member. Simultaneous multi-line electrophysiological evaluation revealed hyperactivity of the simple-variant *CNTN5*^−/+^ iPSC-derived glutamatergic neurons in two independent genetic backgrounds. Moreover, isogenic-MEA and patch-clamp recordings confirmed synaptic hyperactivity of iPSC-derived neurons with disruptive mutations in *EHMT2*, also in two different genetic backgrounds.

To increase the modeling scalability of complex genetic disorders such as ASD while optimizing statistical power, several parameters require careful consideration. Given substantial variation in reprogramming and neuronal differentiation efficiencies, sample size is important to control. It was recently proposed that inter-individual variation, i.e., the number of probands with similar genetic variants, is more important to consider than intra-individual variation, i.e., the number of iPSC clones derived from the same individual (Hoffman et al., 2018). Aiming at multi-variant phenotyping, we tested one or two probands per deficient gene, however, we were able to create an isogenic pair in a different genetic background for the two highly relevant genes, i.e., *CNTN5* (Lionel et al., 2011, Mercati et al., 2017, van Daalen et al., 2011) and *EHMT2* (Deimling et al., 2017, Kleefstra et al., 2005, Zylicz et al., 2015), thereby controlling inter-individual variation. We derived two independent iPSC clones per participants to regulate intra-individual variation. Another important parameter to consider is the cellular homogeneity of neuronal cultures. We preferred to use the NEUROG2 system over classic dual-SMAD inhibition protocols because we believe it represents an advantage in terms of cellular homogeneity. It is also much faster than classic protocols and produces much higher proportion of glutamatergic neurons that can be studied for ASD (Canitano and Pallagrosi, 2017, Habela et al., 2016) or other neurological disorders (Lin et al., 2018).

*NRXN1* (Neurexin 1) is a cell-surface receptor involved in synapse formation and neurotransmission, and was associated with neurodevelopmental disorders with variable penetrance (Tammimies et al., 2015, Woodbury-Smith et al., 2017b). In a mouse knockout model, *Nrxn1* was not essential for excitatory synaptic transmission in cortical neurons, but a significant decrease in mEPSC frequency was observed in a human conditional knockout cell model without change in synapse numbers (Pak et al., 2015). We expected a decreased weighted MFR in **Figure S2A** since our 430-kb deletion disrupts most of the gene and its isoforms, however, we did not detect any significant MEA recordings. Nonetheless, in this instance, no corresponding familial control was available. *DLGAP2* (Discs Large Associated Protein 2) is a major scaffold protein in the post-synaptic density involved in synapse organization and neuronal signaling, and was previously associated with ASD (Marshall et al., 2008). In a mouse model, *Dlgap2* knockout was responsible for a reduction in GluR1 expression and neuronal spine density in the cerebral cortex, and a decreased mEPSC peak amplitude (Jiang-Xie et al., 2014). Our model involved a partial duplication of *DLGAP2* instead of a deletion and no significant results were obtained in our MEA settings, suggesting that this variant had no impact on DLGAP2 protein levels or that the impact is not damaging. *CAPRIN1* (Cell Cycle Associated Protein 1) is an RNA-binding protein involved in proliferation, migration and synaptic plasticity in neurons, and a nonsense variant was identified in a family with ASD (Jiang et al., 2013). A significant reduction in dendrite length and spine density was measured in *Caprin1* knockdown mouse cultured cortical neurons (Shiina and Nakayama, 2014). In neurons from *Caprin1*-/-mice, the frequency and amplitude of mEPSCs was not altered, but the halfwidth of mEPSCs was significantly decreased (Shiina et al., 2010). In addition, a reduced GluR1 expression was detected in primary cortical neurons from *Caprin1*-/-mice (Ohashi et al., 2016). MEA experiments did not reveal any significant differences with our *CAPRIN1*^+/-^ proband-derived neurons, suggesting that neuronal activity might not be strongly affected by the loss of only one *CAPRIN1* allele.

*ANOS1* (Anosmin 1) is a glycoprotein of the extracellular matrix including four consecutive fibronectin type III domains. Loss-of-function variants in *ANOS1* were shown to cause the Kallmann syndrome, which is characterized by congenital hypogonadotropic hypogonadism associated with anosmia, delayed puberty and infertility (Dode and Hardelin, 2009). Defects in the migration of gonadotropin-releasing hormone (GnRH) neurons were observed during embryonic development, as well as morphological changes in the basal forebrain cortex (Manara et al., 2014). ANOS1 expression was detected in projecting neurons and interneurons in the cerebral cortex throughout layers II to VI in rat brains at postnatal day 0 to 15 (Clemente et al., 2008). In human, a proband carrying the nonsense variant R423X in *ANOS1*, and presenting clinical hypogonadotropic hypogonadism, was also diagnosed with ASD (Jiang et al., 2013), suggesting a link between *ANOS1* and ASD. Despite the absence of significant MEA results, neuronal membrane current defects were validated using patch-clamp recordings in two unrelated isogenic pairs (Figure 4D). These results indicate that glutamatergic neuron activity is also influenced by *ANOS1*, which represents a risk gene for ASD. Future studies on iPSC-NEUROG2 neurons will benefit from both MEA and patch-clamp electrophysiology approaches, to uncover subtle deficits.

*CNTN5* (Contactin 5) is an immunoglobulin cell adhesion molecule, with four fibronectin type III domains, involved in neurite outgrowth and axon connection in cortical neurons, and was associated with ASD (van Daalen et al., 2011). Different CNVs affecting *CNTN5* have been associated with ASD and ADHD, with increased occurrence of hyperacusis (Lionel et al., 2011, Mercati et al., 2017). Expression of Cntn5 mRNA was detected in cortical layer IV in mice (Kleijer et al., 2018). Moreover, Cntn5 is expressed in glutamatergic neurons of the central auditory system in mice during the first postnatal week, and knockout of Cntn5 expression altered synapse formation (Toyoshima et al., 2009). The molecular mechanisms through which heterozygous loss of *CNTN5* increases neuronal activity *in vitro* (Figure 3A-B) remains to be elucidated. Gene editing of the 676-kb deletion, as found in lines 27H and 27N (Figure 1D and Table 1), to obtain isogenic controls may be challenging due to the size, but this approach might eventually be applied.

Using a yeast complementation assay (**Figure S4**), we estimated the missense variant E78K in UBE3I not responsible for the electrophysiological phenotypes observed in participant 6-0393-003 (Figure 5A-B). These results prompted us to investigate further on the potential role of the frameshift variant K1164Nfs in *EHMT2*. EHMT2 (G9a) is a histone methyltransferase (HMTase) that forms a complex with EHMT1 (GLP) to catalyze mono- and dimethylation of lysine 9 on histone H3 (H3K9me1/2) (Rice et al., 2003). Of note, EHMT1 protein sequence is highly similar to EHMT2 (Deimling et al., 2017). Actually, EHMT1 haploinsufficiency is involved in intellectual disability (ID) and ASD as part of the Kleefstra syndrome (Kleefstra et al., 2005). EHMT2 represses pluripotency genes in embryonic stem cells (Zylicz et al., 2015) and potentially acts as both repressor of neural progenitor genes and activator of neuronal differentiation genes in early neurodevelopment (Deimling et al., 2017). The impact of the single base deletion in *EHMT2* (K1164Nfs) on the protein function remains to be determined (see Table 1 for details). The frameshift is computationally predicted to extend the protein rather than truncating it, by utilizing sequence in the 3’UTR. However, it is located exactly at the beginning of the post-SET domain, i.e., at position 1164 of EHMT2. The resulting change in the downstream protein sequence completely disrupts three conserved cysteine residues in the post-SET domain that normally form a zinc-binding site with a fourth conserved cysteine close to the SET domain (Zhang et al., 2003). Since these three conserved cysteine residues are essential for HMTase activity, as replacement with serine abolished HMTase activity (Zhang et al., 2002), we suspect that this HMTase activity of EHMT2 is defective in our mutant glutamatergic neurons and potentially related to the observed hyperactivity (Figure 5). Upon further validation experiment using a CRISPR-derived isogenic system and an unrelated genetic background (Figure 5D), we propose that *EHMT2* impacts the synaptic function of glutamatergic neurons through H3K9me1/2 catalyzing ability. Further experiments might clarify this possibility, such as CRISPR-correction of the K1164Nfs point mutation in lines 38B and 38E to obtain isogenic controls.

Overall, this study highlights a way to improve the scalability of testing multiple iPSC-derived neuronal lines with various ASD-risk variants from diverse probands. Furthermore, our work demonstrates that for future studies to capture and characterize the electrophysiological impact of ASD variants on human iPSC-NEUROG2 neurons, it is most beneficial to include both MEA and patch-clamp experiments, across multiple time points. This work also revealed that inactivation of at least one allele of *CNTN5* or *EHMT2* intensifies significantly excitatory neuron synaptic activity *in vitro*. Such phenotype offers the possibility to implement NEUROG2-based high-throughput drug screening strategies (Cheng et al., 2017) combining MEA (Tukker et al., 2018) and lines 38B/38E for instance, to discover molecules that will compensate for neuronal hyperactivity.

## Materials and Methods

### Skin Fibroblasts Culture

Under the approval of the Canadian Institutes of Health Research Stem Cell Oversight Committee, all iPSCs were generated from dermal fibroblasts or CD34+ blood cells. Skin-punch biopsies were obtained from the upper back area by a clinician at The Hospital for Sick Children. Samples were immersed in 14 ml of ice-cold Alpha-MEM (Wisent Bioproducts) supplemented with penicillin 100 Units/ml and streptomycin 100 μg/ml (ThermoFisher), and transferred immediately to the laboratory at The Centre for Applied Genomics (TCAG). Each biopsy was cut into ~1mm^3^ pieces with disposable scalpel in a 60-mm dish. 5 ml of collagenase 1 mg/ml (Sigma, Canada) was added and the dish was placed in 37°C incubator for 1:45 hours. Skin pieces and collagenase were then transferred to a 15-ml tube, and centrifuged at 300 g for 10 minutes. Supernatant was removed, 5 ml of trypsin 0.05%/EDTA 0.53 mM (Wisent Bioproducts) was added, and the mix was pipetted up and down several times to break up tissue and placed in 37°C incubator for 30 minutes. After incubation, the mix was centrifuged at 300 g for 10 minutes, and supernatant was removed leaving 1 ml. The pellet was pipetted up and down vigorously to break to the pieces without creating bubbles. The mix was transferred in a T-12.5 flask along with 5 ml of Alpha-MEM, 15% Fetal Bovine Serum (FBS; Wisent Bioproducts), penicillin 100 Units/ml and streptomycin 100 μg/ml (ThermoFisher), and placed in 37°C incubator for about a week until 100% confluence. Cultured cells were fed every 5-7 days if not confluent. Once confluent, cells were passed into three 100 mm dishes to expand, and frozen in liquid nitrogen.

### Reprogramming Fibroblasts Using Integrative Virus

Reprogramming of skin fibroblasts was performed using retroviral and lentiviral vectors. Retroviral vectors encoding *POU5F1*, *SOX2*, *KLF4*, *MYC*, and lentiviral vectors encoding the pluripotency reporter EOS-GFP/Puro^R^ were used and obtained as described (Hotta et al., 2009).

### Reprogramming Fibroblasts Using Non-Integrative Sendai Virus

Reprogramming of fibroblasts via Sendai virus was performed at the Centre for Commercialization of Regenerative Medicine (CCRM) using CytoTune™-iPS 2.0 Sendai Reprogramming Kit (ThermoFisher). Fibroblasts were cultured in fibroblast expansion media (Advanced DMEM; 10% FBS; 1X L-Glutamine; 1X pen/strep – Thermo Fisher). The desired number of wells for reprogramming from a 24-well plate was coated with 0.1% gelatin. Fibroblasts were dissociated using Trypsin (ThermoFisher) and allowed to settle overnight. Virus multiplicity of infection (MOI) was calculated and viruses combined according to number of cells available for reprogramming and manufacturer’s protocol. 24 hours after transduction, media was changed to wash away viruses. Media was additionally changed on day 3 and 5 after transduction. 6 days after transduction, 6-well plates were coated with Matrigel™(Corning). Cells were removed from the 24-well plate using Accutase (ThermoFisher) and plated on Matrigel in expansion media. 24 hours later, media was replaced with E7 media (StemCellTechnologies). Cells were monitored and fed daily with E7. Once colonies were of an adequate size and morphology to pick, individual colonies were picked and plated into E8 media (StemCellTechnologies). Clones growing well were further expanded and characterized using standard assays for pluripotency, karyotyping, genotyping and mycoplasma testing. Directed differentiation was performed using kits for definitive endoderm, neural and cardiac lineages (all ThermoFisher).

### Peripheral Blood Mononuclear Cells (PBMCs) Isolation from Peripheral Blood and Enrichment of CD34+ Cells

Whole peripheral blood was processed at CCRM using Lymphoprep™ (StemCellTechnologies) in a SepMate™ tube (StemCellTechnologies) according to manufacturer’s instructions. The sample was centrifuged (10 min at 1200 g).

The top layer containing PBMCs was collected and mixed with 10 mL of the PBS/FBS mixture and centrifuged (8 min at 300 g). The PBMC’s collected at the bottom of the tube were washed, counted and resuspended in PBS/FBS mixture. CD34+ cells were then isolated using the Human Whole Blood/Buffy Coat CD34+ Selection kit according to manufacturer’s instructions (StemCellTechnologies). Isolated cells were expanded in StemSpan SFEM II media (StemCellTechnologies) and StemSpan CD34+ Expansion Supplements (StemCellTechnologies) prior to reprogramming.

### Reprogramming PBMC Using Non-Integrative Sendai Virus

Reprogramming of CD34+ PBMCs was performed at CCRM using CytoTune™-iPS 2.0 Sendai Reprogramming Kit. Expanded cells were spun down and resuspended in StemSpan SFEM II media and StemSpan CD34+ Expansion Supplements, and placed in a single well of a 24-well dish. Virus MOI was calculated and viruses combined according to number of cells available for reprogramming and manufacturer’s protocol. The virus mixture was added to cells, and washed off 24 hours after infection. 48 hours after viral delivery, cells were plated in 6-well plates in SFII and transitioned to ReproTESR for the duration of reprogramming. Once colonies were of an adequate size and morphology to pick, individual colonies were picked and plated into E8. Clones growing well were further expanded and characterized as explained above.

### iPSCs Maintenance

All iPSC lines were maintained on matrigel (Corning) coating, with complete media change every day in mTeSR™ (StemCellTechnologies). ReLeSR™ (StemCellTechnologies) was used for passaging. Accutase™ (InnovativeCellTechnologies) and 10 μM Rho-associated kinase (ROCK) inhibitor (Y-27632; StemCellTechnologies) were used for single-cell dissociation purposes.

### Gene Editing

For point mutation correction in 18C line, we used the type II CRISPR/Cas9 double-nicking (Cas9D10A) system with two guide RNA (gRNAs) to reduce off-target activity. We devised the gRNA sequences using tools available at http://crispr.mit.edu/. We designed a HDR-based method using a synthesized single-stranded oligonucleotide (ssODN) template to replace the point mutation with the reference nucleotide. To prevent damage to the correct sequence, a silent mutation was introduced in the ssODN close to the proto-adjacent motif (PAM) of the reverse gRNA (gRNA-), which commands Cas9D10A to nick the plus strand, given that ssODN was synthesized as plus strand. All the CRISPR machinery was introduced into iPSC by nucleofection. Screening for correction of the appropriate base pair was based on absolute quantification of allele frequency using droplet digital PCR (ddPCR). Enrichment of corrected cells was obtained through sib-selection step cultures in 96-well plate format, as adapted from (Miyaoka et al., 2014), until a well containing 100% of corrected alleles was identified. For insertion of premature stop codon in 19-2 cells, ribonucleoprotein (RNP) complex was used as a vector to deliver the CRISPR machinery, along with one sgRNA and Cas9 nuclease, for each target gene. Design of sgRNA and ssODN for HDR, nucleofection and isolation of edited lines were described (Deneault et al., 2018).

### Lentivirus Production

7.5×10^6^ HEK293T cells were seeded in a T-75 flask, grown in 10% fetal bovine serum in DMEM (Gibco). The next day, cells were transfected using Lipofectamine 2000 with plasmids for gag-pol (10 μg), rev (10 μg), VSV-G (5 μg), and the target constructs FUW-TetO-Ng2-P2A-EGFP-T2A-puromycin or FUW-rtTA (15 μg; gift from T.C. Südhof laboratory) (Zhang et al., 2013). Next day, the media was changed. The day after that, the media was spun down in a high-speed centrifuge at 30,000 g at 4°C for 2 hours. The supernatant was discarded and 50 μl PBS was added to the pellet and left overnight at 4°C. The next day, the solution was triturated, aliquoted and frozen at −80°C.

### Differentiation into Glutamatergic Neurons

5×10^5^ iPSCs/well were seeded in a matrigel-coated 6-well plate in 2 ml of mTeSR supplemented with 10 μM Y-27632. Next day, media in each well was replaced with 2 ml fresh media plus 10 μM Y-27632, 0.8 μg/ml polybrene (Sigma), and the minimal amount of NEUROG2 and rtTA lentiviruses necessary to generate 100% GFP+ cells upon doxycycline induction, depending on prior titration of a given virus batch. The day after, virus-containing media were replaced with fresh mTeSR, and cells were expanded until near-confluency. Newly generated “NEUROG2-iPSCs” were detached using accutase, and seeded in a new matrigel-coated 6-well plate at a density of 5×10^5^ cells per well in 2 ml of mTeSR supplemented with 10 μM Y-27632 (day 0 of differentiation). Next day (day 1), media in each well was changed for 2 ml of CM1 [DMEM-F12 (Gibco), 1x N2 (Gibco), 1x NEAA (Gibco), 1x pen/strep (Gibco), laminin (1 μg/ml; Sigma), BDNF (10 ng/μl; Peprotech) and GDNF (10 ng/μl; Peprotech) supplemented with fresh doxycycline hyclate (2 μg/ml; Sigma) and 10 μM Y-27632. The day after (day 2), media was replaced with 2 ml of CM2 [Neurobasal media (Gibco), 1x B27 (Gibco), 1x glutamax (Gibco), 1x pen/strep, laminin (1 μg/ml), BDNF (10 ng/μl) and GDNF (10 ng/μl)] supplemented with fresh doxycycline hyclate (2 μg/ml) and puromycin (5 μg/ml for 19-2-derived cells, and 2 μg/ml for 50B-derived cells; Sigma). Media was replaced with CM2 supplemented with fresh doxycycline hyclate (2 μg/ml). The same media change was repeated at day 4. At day 6, media was replaced with CM2 supplemented with fresh doxycycline hyclate (2 μg/ml) and araC (10 μM; Sigma). Two days later, these day 8 post-NEUROG2-induction (PNI) neurons were detached using accutase and ready to seed for subsequent experiments, as described below.

### Multi-electrode array (MEA)

48-well opaque-bottom MEA plates (Axion Biosystems) were coated with filter-sterilized 0.1% polyethyleneimine solution in borate buffer pH 8.4 for 1 hour at room temperature, washed four times with water, and dried overnight. 120,000 “day8-dox” neurons/well were seeded in 250 μl CM2 media. The day after, 5,000 mouse astrocytes/well were seeded on top of neurons in 50 μl/well CM2 media. Astrocytes were prepared from postnatal day 1 CD-1 mice as described (Kim and Magrane, 2011). Media was half-changed once a week with CM2 media. Every week post-seeding, the electrical activity of the MEA plates was recorded using the Axion Maestro MEA reader (Axion Biosystems). The heater control was set to warm up the reader at 37°C. Each plate was first incubated for 5 minutes on the pre-warmed reader, then real-time spontaneous neural activity was recorded for 5 minutes using AxIS 2.0 software (Axion Biosystems). A bandpass filter from 200 Hz to 3 kHz was applied. Spikes were detected using a threshold of 6 times the standard deviation of noise signal on electrodes.

Offline advanced metrics were re-recorded and analysed using Axion Biosystems Neural Metric Tool. An electrode was considered active if at least 5 spikes were detected per minute. Single electrode bursts were identified as a minimum of 5 spikes with a maximum interspike interval (ISI) of 100 milliseconds. Network bursts were identified as a minimum of 10 spikes with a maximum ISI of 100 milliseconds covered by at least 25% of electrodes in each well. No non-active well was excluded in the analysis. After the last reading, each well was treated with three synaptic antagonists: GABA_A_ receptor antagonist picrotoxin (PTX; Sigma) at 100 μM, AMPA receptor antagonist 6-cyano-7-nitroquinoxaline-2,3-dion (CNQX; Sigma) at 60 μM, and sodium ion channel antagonist tetrodotoxin (TTX; Alomone labs) at 1 μM. The plates were recorded consecutively, 5-10 minutes after addition of the antagonists. A 60-minute recovery period was allowed in the incubator at 37°C between each antagonist treatment and plate recording.

### Patch-Clamp Recordings

Day 3 PNI neurons were replated at a density of 100 000/well of a poly-ornithin/laminin coated coverslips in a 24-well plate with CM2 media. On day 4, 50 000 mouse astrocytes were added to the plates and cultured until day 21-28 PNI for recording. At day 10, CM2 was supplemented with 2.5% FBS in accordance with (Zhang et al., 2013). Whole-cell recordings (BX51WI; Olympus) were performed at room temperature using an Axoclamp 700B amplifier (Molecular Devices) from borosilicate patch electrodes (P-97 puller; Sutter Instruments) containing a potassium-based intracellular solution (in mM): 123 K-gluconate, 10 KCL, 10 HEPES; 1 EGTA, 2 MgCl_2_, 0.1 CaCl_2_, 1 Mg_-_ATP, and 0.2 Na_4_GTP (pH 7.2). 0.06% sulpharhodamine dye was added to select neurons for visual confirmation of multipolar neurons. Composition of extracellular solution was (in mM): 140 NaCl, 2.5 KCl, 1 1.25 NaH_2_PO_4_, 1 MgCl_2_, 10 glucose, and 2 CaCl_2_ (pH 7.4). Whole cell recordings were clamped at −70 mV using Clampex 10.6 (Molecular Devices), corrected for a calculated −10 mV junction potential and analyzed using the Template Search function from Clampfit 10.6 (Molecular Devices). Following initial breakthrough and current stabilization in voltage clamp, the cell was switched to current clamp to monitor initial spiking activity and record the membrane potential (cc=0, ~1 min post-breakthrough). Bias current was applied to bring the cell to ~70 mV whereby increasing 5 pA current steps were applied (starting at −,20 pA) to generate the whole cell resistance and to elicit action potentials. Data were digitized at 10 kHz and low-pass filtered at 2 kHz. Inward and outward currents were recorded in whole-cell voltage clamp in response to consecutive 10 mV steps from −90 mV to +40 mV.

### Yeast Complementation Assay

The method for the yeast complementation assay was described previously (Sun et al., 2016).

### Antibodies and Western Blotting

Cells were washed in ice-cold PBS and total protein was extracted in RIPA supplemented with proteinase inhibitor cocktail, and homogenized. Equivalent protein mass was loaded on gradient SDS-PAGE (4-12%) and transferred to Nitrocellulose membrane Hybond ECL (GE HealthCare). Primary antibodies used were rabbit anti-CNTN5 (Novus, NBP1-83243) and rabbit anti-EHMT2/G9A (Abcam, ab185050). HRP-conjugated secondary antibodies (Invitrogen) were used and the membranes were developed with SuperSignal West Pico Chemiluminescent Substrate (Pierce). Images acquired using ChemiDoc MP (BioRad) and quantified using software Imagelab v4.1 (BioRad). Western Blots were repeated at least twice for each biological replicate.

### Mycoplasma Testing

All cell lines were regularly tested for presence of mycoplasma using a standard method (Otto, 1996).

## Acknowledgements

We thank the Autism Speaks MSSNG project for genomic data and linking to consented families. We also thank Melissa Carter, Wendy Roberts, Brian Chung and Rosanna Weksberg for obtaining skin biopsies and blood work, and the families for volunteering. We also thank Tara Paton, Guillermo Casallo, Barbara Kellam, Ny Hoang and Sylvia Lamoureux for technical help; T.C. Südhof for the NEUROG2/rtTA lentiviral constructs.

## References

Beltrao-braga, P. C. & Muotri, A. R. 2017. Modeling autism spectrum disorders with human neurons. Brain Res, 1656, 49-54.

Betancur, C. 2011. Etiological heterogeneity in autism spectrum disorders: more than 100 genetic and genomic disorders and still counting. Brain Res, 1380, 42-77.

Bishop, S. L., Farmer, C., Bal, V., Robinson, E. B., Willsey, A. J., Werling, D. M., Havdahl, K. A., Sanders, S. J. & Thurm, A. 2017. Identification of Developmental and Behavioral Markers Associated With Genetic Abnormalities in Autism Spectrum Disorder. Am J Psychiatry, 174, 576-585.

Bourgeron, T. 2015. From the genetic architecture to synaptic plasticity in autism spectrum disorder. Nat Rev Neurosci, 16, 551-63.

Brandler, W. M., Antaki, D., Gujral, M., Kleiber, M. L., Whitney, J., Maile, M. S., Hong, O., Chapman, T. R., Tan, S., Tandon, P., Pang, T., Tang, S. C., Vaux, K. K., Yang, Y., Harrington, E., Juul, S., Turner, D. J., Thiruvahindrapuram, B., Kaur, G., Wang, Z., Kingsmore, S. F., Gleeson, J. G., Bisson, D., Kakaradov, B., Telenti, A., Venter, J. C., Corominas, R., Toma, C., Cormand, B., Rueda, I., Guijarro, S., Messer, K. S., Nievergelt, C. M., Arranz, M. J., Courchesne, E., Pierce, K., Muotri, A. R., Iakoucheva, L. M., Hervas, A., Scherer, S. W., Corsello, C. & Sebat, J. 2018. Paternally inherited cis-regulatory structural variants are associated with autism. Science, 360, 327-331.

Canitano, R. & Pallagrosi, M. 2017. Autism Spectrum Disorders and Schizophrenia Spectrum Disorders: Excitation/Inhibition Imbalance and Developmental Trajectories. Front Psychiatry, 8, 69.

Carter, M. T. & Scherer, S. W. 2013. Autism spectrum disorder in the genetics clinic: a review. Clin Genet, 83, 399-407.

Cheng, C., Fass, D. M., Folz-Donahue, K., Macdonald, M. E. & Haggarty, S. J. 2017. Highly Expandable Human iPS Cell-Derived Neural Progenitor Cells (NPC) and Neurons for Central Nervous System Disease Modeling and High-Throughput Screening. Curr Protoc Hum Genet, 92, 21 8 1-21 8 21.

Clemente, D., Esteban, P. F., Del Valle, I., Bribian, A., Soussi-Yanicostas, N., Silva, A. & De Castro, F. 2008. Expression pattern of Anosmin-1 during pre- and postnatal rat brain development. Dev Dyn, 237, 2518-28.

De Rubeis, S., He, X., Goldberg, A. P., Poultney, C. S., Samocha, K., Cicek, A. E., Kou, Y., Liu, L., Fromer, M., Walker, S., Singh, T., Klei, L., Kosmicki, J., Shih-Chen, F., Aleksic, B., Biscaldi, M., Bolton, P. F., Brownfeld, J. M., Cai, J., Campbell, N. G., Carracedo, A., Chahrour, M. H., Chiocchetti, A. G., Coon, H., Crawford, E. L., Curran, S. R., Dawson, G., Duketis, E., Fernandez, B. A., Gallagher, L., Geller, E., Guter, S. J., Hill, R. S., Ionita-Laza, J., Jimenz Gonzalez, P., Kilpinen, H., Klauck, S. M., Kolevzon, A., Lee, I., Lei, I., Lei, J., Lehtimaki, T., Lin, C. F., Ma’ayan, A., Marshall, C. R., Mcinnes, A. L., Neale, B., Owen, M. J., Ozaki, N., Parellada, M., Parr, J. R., Purcell, S., Puura, K., Rajagopalan, D., Rehnstrom, K., Reichenberg, A., Sabo, A., Sachse, M., Sanders, S. J., Schafer, C., Schulte-Ruther, M., Skuse, D., Stevens, C., Szatmari, P., Tammimies, K., Valladares, O., Voran, A., Li-San, W., Weiss, L. A., Willsey, A. J., Yu, T. W., Yuen, R. K., Study, D. D. D., Homozygosity Mapping Collaborative For, A., Consortium, U. K., Cook, E. H., Freitag, C. M., Gill, M., Hultman, C. M., Lehner, T., Palotie, A., Schellenberg, G. D., Sklar, P., State, M. W., Sutcliffe, J. S., Walsh, C. A., Scherer, S. W., Zwick, M. E., Barett, J. C., Cutler, D. J., Roeder, K., Devlin, B., Daly, M. J. & Buxbaum, J. D. 2014. Synaptic, transcriptional and chromatin genes disrupted in autism. Nature, 515, 209-15.

Deimling, S. J., Olsen, J. B. & Tropepe, V. 2017. The expanding role of the Ehmt2/G9a complex in neurodevelopment. Neurogenesis (Austin), 4, e1316888.

Deneault, E., White, S. H., Rodrigues, D., Ross, P. J., Faheem, M., Zaslavsky, K., Wang, Z., Alexandrova, R., Pellecchia, G., Wei, W., Piekna, A., Kaur, G., Howe, J. L., Kwan, V., Thiruvahindrapuram, B., Walker, S., Pasceri, P., Merico, D., Yuen, R. C. K., Singh, K. K., Ellis, J. & Scherer, S. W. 2018. Complete Disruption of Autism-Susceptibility Genes by Gene-Editing Predominantly Reduces Functional Connectivity of Isogenic Human Neurons. bioRxiv preprint server, DOI:10.1101/344234.

Devlin, B. & Scherer, S. W. 2012. Genetic architecture in autism spectrum disorder. Curr Opin Genet Dev, 22, 229-37.

Dode, C. & Hardelin, J. P. 2009. Kallmann syndrome. Eur J Hum Genet, 17, 139-46.

Dolmetsch, R. & Geschwind, D. H. 2011. The human brain in a dish: the promise of iPSC-derived neurons. Cell, 145, 831-4.

DSM-V 2013. Diagnostic and Statistical Manual of Mental Disorders, Fifth Edition. American Psychiatric Association.

Durak, O. & Tsai, L. H. 2014. Human induced pluripotent stem cells: now open to discovery. Cell Stem Cell, 15, 4-6.

Fernandez, B. A. & Scherer, S. W. 2017. Syndromic autism spectrum disorders: moving from a clinically defined to a molecularly defined approach. Dialogues Clin Neurosci, 19, 353-371.

Geschwind, D. H. & State, M. W. 2015. Gene hunting in autism spectrum disorder: on the path to precision medicine. Lancet Neurol, 14, 1109-20.

Gilman, S. R., Iossifov, I., Levy, D., Ronemus, M., Wigler, M. & Vitkup, D. 2011. Rare de novo variants associated with autism implicate a large functional network of genes involved in formation and function of synapses. Neuron, 70, 898-907.

Habela, C. W., Song, H. & Ming, G. L. 2016. Modeling synaptogenesis in schizophrenia and autism using human iPSC derived neurons. Mol Cell Neurosci, 73, 52-62.

Ho, S. M., Hartley, B. J., Tcw, J., Beaumont, M., Stafford, K., Slesinger, P. A. & Brennand, K. J. 2016. Rapid Ngn2-induction of excitatory neurons from hiPSC-derived neural progenitor cells. Methods, 101, 113-24.

Hoffman, G. E., Schrode, N., Flaherty, E. & Brennand, K. J. 2018. New considerations for hiPSC-based models of neuropsychiatric disorders. Mol Psychiatry.

Hotta, A., Cheung, A. Y., Farra, N., Vijayaragavan, K., Seguin, C. A., Draper, J. S., Pasceri, P., Maksakova, I. A., Mager, D. L., Rossant, J., Bhatia, M. & Ellis, J. 2009. Isolation of human iPS cells using EOS lentiviral vectors to select for pluripotency. Nat Methods, 6, 370-6.

Jiang, Y. H., Yuen, R. K., Jin, X., Wang, M., Chen, N., Wu, X., Ju, J., Mei, J., Shi, Y., He, M., Wang, G., Liang, J., Wang, Z., Cao, D., Carter, M. T., Chrysler, C., Drmic, I. E., Howe, J. L., Lau, L., Marshall, C. R., Merico, D., Nalpathamkalam, T., Thiruvahindrapuram, B., Thompson, A., Uddin, M., Walker, S., Luo, J., Anagnostou, E., Zwaigenbaum, L., Ring, R. H., Wang, J., Lajonchere, C., Wang, J., Shih, A., Szatmari, P., Yang, H., Dawson, G., Li, Y. & Scherer, S. W. 2013. Detection of clinically relevant genetic variants in autism spectrum disorder by whole-genome sequencing. Am J Hum Genet, 93, 249-63.

Jiang-Xie, L. F., Liao, H. M., Chen, C. H., Chen, Y. T., Ho, S. Y., Lu, D. H., Lee, L. J., Liou, H. H., Fu, W. M. & Gau, S. S. 2014. Autism-associated gene Dlgap2 mutant mice demonstrate exacerbated aggressive behaviors and orbitofrontal cortex deficits. Mol Autism, 5, 32.

Jinek, M., Chylinski, K., Fonfara, I., Hauer, M., Doudna, J. A. & Charpentier, E. 2012. A programmable dual-RNA-guided DNA endonuclease in adaptive bacterial immunity. Science, 337, 816-21.

Jones, R. M. & Lord, C. 2013. Diagnosing autism in neurobiological research studies. Behav Brain Res, 251, 113-24.

Karmacharya, R. & Haggarty, S. J. 2016. Stem cell models of neuropsychiatric disorders. Mol Cell Neurosci, 73, 1-2.

Kim, H. J. & Magrane, J. 2011. Isolation and culture of neurons and astrocytes from the mouse brain cortex. Methods Mol Biol, 793, 63-75.

Kleefstra, T., Smidt, M., Banning, M. J., Oudakker, A. R., Van Esch, H., De Brouwer, A. P., Nillesen, W., Sistermans, E. A., Hamel, B. C., De Bruijn, D., Fryns, J. P., Yntema, H. G., Brunner, H. G., De Vries, B. B. & Van Bokhoven, H. 2005. Disruption of the gene Euchromatin Histone Methyl Transferase1 (Eu-HMTase1) is associated with the 9q34 subtelomeric deletion syndrome. J Med Genet, 42, 299-306.

Kleijer, K. T. E., Van Nieuwenhuize, D., Spierenburg, H. A., Gregorio-Jordan, S., Kas, M. J. H. & Burbach, J. P. H. 2018. Structural abnormalities in the primary somatosensory cortex and a normal behavioral profile in Contactin-5 deficient mice. Cell Adh Migr, 12, 5-18.

Lin, Y. T., Seo, J., Gao, F., Feldman, H. M., Wen, H. L., Penney, J., Cam, H. P., Gjoneska, E., Raja, W. K., Cheng, J., Rueda, R., Kritskiy, O., Abdurrob, F., Peng, Z., Milo, B., Yu, C. J., Elmsaouri, S., Dey, D., Ko, T., Yankner, B. A. & Tsai, L. H. 2018. APOE4 Causes Widespread Molecular and Cellular Alterations Associated with Alzheimer’s Disease Phenotypes in Human iPSC-Derived Brain Cell Types. Neuron, 98, 1294.

Lionel, A. C., Crosbie, J., Barbosa, N., Goodale, T., Thiruvahindrapuram, B., Rickaby, J., Gazzellone, M., Carson, A. R., Howe, J. L., Wang, Z., Wei, J., Stewart, A. F., Roberts, R., Mcpherson, R., Fiebig, A., Franke, A., Schreiber, S., Zwaigenbaum, L., Fernandez, B. A., Roberts, W., Arnold, P. D., Szatmari, P., Marshall, C. R., Schachar, R. & Scherer, S. W. 2011. Rare copy number variation discovery and cross-disorder comparisons identify risk genes for ADHD. Sci Transl Med, 3, 95ra75.

Mahdi, S., Albertowski, K., Almodayfer, O., Arsenopoulou, V., Carucci, S., Dias, J. C., Khalil, M., Knuppel, A., Langmann, A., Lauritsen, M. B., Da Cunha, G. R., Uchiyama, T., Wolff, N., Selb, M., Granlund, M., De Vries, P. J., Zwaigenbaum, L. & Bolte, S. 2018. An International Clinical Study of Ability and Disability in Autism Spectrum Disorder Using the WHO-ICF Framework. J Autism Dev Disord, 48, 2148-2163.

Malhotra, D. & Sebat, J. 2012. CNVs: harbingers of a rare variant revolution in psychiatric genetics. Cell, 148, 1223-41.

Manara, R., Salvalaggio, A., Favaro, A., Palumbo, V., Citton, V., Elefante, A., Brunetti, A., Di Salle, F., Bonanni, G., Sinisi, A. A. & Kallmann Syndrome Neuroradiological Study, G. 2014. Brain changes in Kallmann syndrome. AJNR Am J Neuroradiol, 35, 1700-6.

Marchetto, M. C., Belinson, H., Tian, Y., Freitas, B. C., Fu, C., Vadodaria, K., Beltrao-Braga, P., Trujillo, C. A., Mendes, A. P. D., Padmanabhan, K., Nunez, Y., Ou, J., Ghosh, H., Wright, R., Brennand, K., Pierce, K., Eichenfield, L., Pramparo, T., Eyler, L., Barnes, C. C., Courchesne, E., Geschwind, D. H., Gage, F. H., Wynshaw-Boris, A. & Muotri, A. R. 2017. Altered proliferation and networks in neural cells derived from idiopathic autistic individuals. Mol Psychiatry, 22, 820-835.

Marshall, C. R., Noor, A., Vincent, J. B., Lionel, A. C., Feuk, L., Skaug, J., Shago, M., Moessner, R., Pinto, D., Ren, Y., Thiruvahindrapduram, B., Fiebig, A., Schreiber, S., Friedman, J., Ketelaars, C. E., Vos, Y. J., Ficicioglu, C., Kirkpatrick, S., Nicolson, R., Sloman, L., Summers, A., Gibbons, C. A., Teebi, A., Chitayat, D., Weksberg, R., Thompson, A., Vardy, C., Crosbie, V., Luscombe, S., Baatjes, R., Zwaigenbaum, L., Roberts, W., Fernandez, B., Szatmari, P. & Scherer, S. W. 2008. Structural variation of chromosomes in autism spectrum disorder. Am J Hum Genet, 82, 477-88.

Mercati, O., Huguet, G., Danckaert, A., Andre-Leroux, G., Maruani, A., Bellinzoni, M., Rolland, T., Gouder, L., Mathieu, A., Buratti, J., Amsellem, F., Benabou, M., Van-Gils, J., Beggiato, A., Konyukh, M., Bourgeois, J. P., Gazzellone, M. J., Yuen, R. K., Walker, S., Delepine, M., Boland, A., Regnault, B., Francois, M., Van Den Abbeele, T., Mosca-Boidron, A. L., Faivre, L., Shimoda, Y., Watanabe, K., Bonneau, D., Rastam, M., Leboyer, M., Scherer, S. W., Gillberg, C., Delorme, R., Cloez-Tayarani, I. & Bourgeron, T. 2017. CNTN6 mutations are risk factors for abnormal auditory sensory perception in autism spectrum disorders. Mol Psychiatry, 22, 625-633.

Miyaoka, Y., Chan, A. H., Judge, L. M., Yoo, J., Huang, M., Nguyen, T. D., Lizarraga, P. P., So, P. L. & Conklin, B. R. 2014. Isolation of single-base genome-edited human iPS cells without antibiotic selection. Nat Methods, 11, 291-3.

Ohashi, R., Takao, K., Miyakawa, T. & Shiina, N. 2016. Comprehensive behavioral analysis of RNG105 (Caprin1) heterozygous mice: Reduced social interaction and attenuated response to novelty. Sci Rep, 6, 20775.

Otto, E. Z., C; Kaloss, M; Del Giudice, Ra; Gardella, R; Mcgarrity, GK. 1996. Quantitative detection of cell culture Mycoplasmas by a one step polymerase chain reaction method. Methods in Cell Science, 18, 261-268.

Pak, C., Danko, T., Zhang, Y., Aoto, J., Anderson, G., Maxeiner, S., Yi, F., Wernig, M. & Sudhof, T. C. 2015. Human Neuropsychiatric Disease Modeling using Conditional Deletion Reveals Synaptic Transmission Defects Caused by Heterozygous Mutations in NRXN1. Cell Stem Cell, 17, 316-28.

Pinto, D., Delaby, E., Merico, D., Barbosa, M., Merikangas, A., Klei, L., Thiruvahindrapuram, B., Xu, X., Ziman, R., Wang, Z., Vorstman, J. A., Thompson, A., Regan, R., Pilorge, M., Pellecchia, G., Pagnamenta, A. T., Oliveira, B., Marshall, C. R., Magalhaes, T. R., Lowe, J. K., Howe, J. L., Griswold, A. J., Gilbert, J., Duketis, E., Dombroski, B. A., De Jonge, M. V., Cuccaro, M., Crawford, E. L., Correia, C. T., Conroy, J., Conceicao, I. C., Chiocchetti, A. G., Casey, J. P., Cai, G., Cabrol, C., Bolshakova, N., Bacchelli, E., Anney, R., Gallinger, S., Cotterchio, M., Casey, G., Zwaigenbaum, L., Wittemeyer, K., Wing, K., Wallace, S., Van Engeland, H., Tryfon, A., Thomson, S., Soorya, L., Roge, B., Roberts, W., Poustka, F., Mouga, S., Minshew, N., Mcinnes, L. A., Mcgrew, S. G., Lord, C., Leboyer, M., Le Couteur, A. S., Kolevzon, A., Jimenez Gonzalez, P., Jacob, S., Holt, R., Guter, S., Green, J., Green, A., Gillberg, C., Fernandez, B. A., Duque, F., Delorme, R., Dawson, G., Chaste, P., Cafe, C., Brennan, S., Bourgeron, T., Bolton, P. F., Bolte, S., Bernier, R., Baird, G., Bailey, A. J., Anagnostou, E., Almeida, J., Wijsman, E. M., Vieland, V. J., Vicente, A. M., Schellenberg, G. D., Pericak-Vance, M., Paterson, A. D., Parr, J. R., Oliveira, G., Nurnberger, J. I., Monaco, A. P., Maestrini, E., Klauck, S. M., Hakonarson, H., Haines, J. L., Geschwind, D. H., Freitag, C. M., Folstein, S. E., Ennis, S., et al. 2014. Convergence of genes and cellular pathways dysregulated in autism spectrum disorders. Am J Hum Genet, 94, 677-94.

Powell, S. K., Gregory, J., Akbarian, S. & Brennand, K. J. 2017. Application of CRISPR/Cas9 to the study of brain development and neuropsychiatric disease. Mol Cell Neurosci, 82, 157-166.

Ran, F. A., Hsu, P. D., Lin, C. Y., Gootenberg, J. S., Konermann, S., Trevino, A. E., Scott, D. A., Inoue, A., Matoba, S., Zhang, Y. & Zhang, F. 2013. Double nicking by RNA-guided CRISPR Cas9 for enhanced genome editing specificity. Cell, 154, 1380-9.

Rice, J. C., Briggs, S. D., Ueberheide, B., Barber, C. M., Shabanowitz, J., Hunt, D. F., Shinkai, Y. & Allis, C. D. 2003. Histone methyltransferases direct different degrees of methylation to define distinct chromatin domains. Mol Cell, 12, 1591-8.

Ronald, A. & Hoekstra, R. A. 2011. Autism spectrum disorders and autistic traits: a decade of new twin studies. Am J Med Genet B Neuropsychiatr Genet, 156B, 255-74.

Roybon, L., Mastracci, T. L., Ribeiro, D., Sussel, L., Brundin, P. & Li, J. Y. 2010. GABAergic differentiation induced by Mash1 is compromised by the bHLH proteins Neurogenin2, NeuroD1, and NeuroD2. Cereb Cortex, 20, 1234-44.

Sahin, M. & Sur, M. 2015. Genes, circuits, and precision therapies for autism and related neurodevelopmental disorders. Science, 350.

Sanders, S. J., He, X., Willsey, A. J., Ercan-Sencicek, A. G., Samocha, K. E., Cicek, A. E., Murtha, M. T., Bal, V. H., Bishop, S. L., Dong, S., Goldberg, A. P., Jinlu, C., Keaney, J. F., 3rd, Klei, L., Mandell, J. D., Moreno-de-luca, D., Poultney, C. S., Robinson, E. B., Smith, L., Solli-Nowlan, T., Su, M. Y., Teran, N. A., Walker, M. F., Werling, D. M., Beaudet, A. L., Cantor, R. M., Fombonne, E., Geschwind, D. H., Grice, D. E., Lord, C., Lowe, J. K., Mane, S. M., Martin, D. M., Morrow, E. M., Talkowski, M. E., Sutcliffe, J. S., Walsh, C. A., Yu, T. W., Autism Sequencing, C., Ledbetter, D. H., Martin, C. L., Cook, E. H., Buxbaum, J. D., Daly, M. J., Devlin, B., Roeder, K. & State, M. W. 2015. Insights into Autism Spectrum Disorder Genomic Architecture and Biology from 71 Risk Loci. Neuron, 87, 1215-1233.

Shiina, N. & Nakayama, K. 2014. RNA granule assembly and disassembly modulated by nuclear factor associated with double-stranded RNA 2 and nuclear factor 45. J Biol Chem, 289, 21163-80.

Shiina, N., Yamaguchi, K. & Tokunaga, M. 2010. RNG105 deficiency impairs the dendritic localization of mRNAs for Na+/K+ ATPase subunit isoforms and leads to the degeneration of neuronal networks. J Neurosci, 30, 12816-30.

Sun, S., Yang, F., Tan, G., Costanzo, M., Oughtred, R., Hirschman, J., Theesfeld, C. L., Bansal, P., Sahni, N., Yi, S., Yu, A., Tyagi, T., Tie, C., Hill, D. E., Vidal, M., Andrews, B. J., Boone, C., Dolinski, K. & Roth, F. P. 2016. An extended set of yeast-based functional assays accurately identifies human disease mutations. Genome Res, 26, 670-80.

Takahashi, K., Tanabe, K., Ohnuki, M., Narita, M., Ichisaka, T., Tomoda, K. & Yamanaka, S. 2007. Induction of pluripotent stem cells from adult human fibroblasts by defined factors. Cell, 131, 861-72.

Tammimies, K., Marshall, C. R., Walker, S., Kaur, G., Thiruvahindrapuram, B., Lionel, A. C., Yuen, R. K., Uddin, M., Roberts, W., Weksberg, R., Woodbury-Smith, M., Zwaigenbaum, L., Anagnostou, E., Wang, Z., Wei, J., Howe, J. L., Gazzellone, M. J., Lau, L., Sung, W. W., Whitten, K., Vardy, C., Crosbie, V., Tsang, B., D’Abate, L., Tong, W. W., Luscombe, S., Doyle, T., Carter, M. T., Szatmari, P., Stuckless, S., Merico, D., Stavropoulos, D. J., Scherer, S. W. & Fernandez, B. A. 2015. Molecular Diagnostic Yield of Chromosomal Microarray Analysis and Whole-Exome Sequencing in Children With Autism Spectrum Disorder. JAMA, 314, 895-903.

Toyoshima, M., Sakurai, K., Shimazaki, K., Takeda, Y., Shimoda, Y. & Watanabe, K. 2009. Deficiency of neural recognition molecule NB-2 affects the development of glutamatergic auditory pathways from the ventral cochlear nucleus to the superior olivary complex in mouse. Dev Biol, 336, 192-200.

Tukker, A. M., Wijnolts, F. M. J., De Groot, A. & Westerink, R. H. S. 2018. Human iPSC-derived neuronal models for in vitro neurotoxicity assessment. Neurotoxicology, 67, 215-225.

Uddin, M., Tammimies, K., Pellecchia, G., Alipanahi, B., Hu, P., Wang, Z., Pinto, D., Lau, L., Nalpathamkalam, T., Marshall, C. R., Blencowe, B. J., Frey, B. J., Merico, D., Yuen, R. K. & Scherer, S. W. 2014. Brain-expressed exons under purifying selection are enriched for de novo mutations in autism spectrum disorder. Nat Genet, 46, 742-7.

Van Daalen, E., Kemner, C., Verbeek, N. E., Van Der Zwaag, B., Dijkhuizen, T., Rump, P., Houben, R., Van ’T Slot, R., De Jonge, M. V., Staal, W. G., Beemer, F. A., Vorstman, J. A., Burbach, J. P., Van Amstel, H. K., Hochstenbach, R., Brilstra, E. H. & Poot, M. 2011. Social Responsiveness Scale-aided analysis of the clinical impact of copy number variations in autism. Neurogenetics, 12, 315-23.

Weiner, D. J., Wigdor, E. M., Ripke, S., Walters, R. K., Kosmicki, J. A., Grove, J., Samocha, K. E., Goldstein, J. I., Okbay, A., Bybjerg-Grauholm, J., Werge, T., Hougaard, D. M., Taylor, J., I, P.-B. A. G., Psychiatric Genomics Consortium Autism, G., Skuse, D., Devlin, B., Anney, R., Sanders, S. J., Bishop, S., Mortensen, P. B., Borglum, A. D., Smith, G. D., Daly, M. J. & Robinson, E. B. 2017. Polygenic transmission disequilibrium confirms that common and rare variation act additively to create risk for autism spectrum disorders. Nat Genet, 49, 978-985.

Winden, K. D., Ebrahimi-Fakhari, D. & Sahin, M. 2018. Abnormal mTOR Activation in Autism. Annu Rev Neurosci, 41, 1-23.

Woodbury-Smith, M., Deneault, E., Yuen, R. K. C., Walker, S., Zarrei, M., Pellecchia, G., Howe, J. L., Hoang, N., Uddin, M., Marshall, C. R., Chrysler, C., Thompson, A., Szatmari, P. & Scherer, S. W. 2017a. Mutations in RAB39B in individuals with intellectual disability, autism spectrum disorder, and macrocephaly. Mol Autism, 8, 59.

Woodbury-Smith, M., Nicolson, R., Zarrei, M., Yuen, R. K. C., Walker, S., Howe, J., Uddin, M., Hoang, N., Buchanan, J. A., Chrysler, C., Thompson, A., Szatmari, P. & Scherer, S. W. 2017b. Variable phenotype expression in a family segregating microdeletions of the NRXN1 and MBD5 autism spectrum disorder susceptibility genes. NPJ Genom Med, 2.

Yi, F., Danko, T., Botelho, S. C., Patzke, C., Pak, C., Wernig, M. & Sudhof, T. C. 2016. Autism-associated SHANK3 haploinsufficiency causes Ih channelopathy in human neurons. Science, 352, aaf2669.

Yoon, K. J., Nguyen, H. N., Ursini, G., Zhang, F., Kim, N. S., Wen, Z., Makri, G., Nauen, D., Shin, J. H., Park, Y., Chung, R., Pekle, E., Zhang, C., Towe, M., Hussaini, S. M., Lee, Y., Rujescu, D., St Clair, D., Kleinman, J. E., Hyde, T. M., Krauss, G., Christian, K. M., Rapoport, J. L., Weinberger, D. R., Song, H. & Ming, G. L. 2014. Modeling a genetic risk for schizophrenia in iPSCs and mice reveals neural stem cell deficits associated with adherens junctions and polarity. Cell Stem Cell, 15, 79-91.

Yu, J., Vodyanik, M. A., Smuga-Otto, K., Antosiewicz-bourget, J., Frane, J. L., Tian, S., Nie, J., Jonsdottir, G. A., Ruotti, V., Stewart, R., Slukvin, II & Thomson, J. A. 2007. Induced pluripotent stem cell lines derived from human somatic cells. Science, 318, 1917-20.

Yuen, R. K., Merico, D., Bookman, M., J, L. H., Thiruvahindrapuram, B., Patel, R. V., Whitney, J., Deflaux, N., Bingham, J., Wang, Z., Pellecchia, G., Buchanan, J. A., Walker, S., Marshall, C. R., Uddin, M., Zarrei, M., Deneault, E., D’abate, L., Chan, A. J., Koyanagi, S., Paton, T., Pereira, S. L., Hoang, N., Engchuan, W., Higginbotham, E. J., Ho, K., Lamoureux, S., Li, W., Macdonald, J. R., Nalpathamkalam, T., Sung, W. W., Tsoi, F. J., Wei, J., Xu, L., Tasse, A. M., Kirby, E., Van Etten, W., Twigger, S., Roberts, W., Drmic, I., Jilderda, S., Modi, B. M., Kellam, B., Szego, M., Cytrynbaum, C., Weksberg, R., Zwaigenbaum, L., Woodbury-Smith, M., Brian, J., Senman, L., Iaboni, A., Doyle-Thomas, K., Thompson, A., Chrysler, C., Leef, J., Savion-Lemieux, T., Smith, I. M., Liu, X., Nicolson, R., Seifer, V., Fedele, A., Cook, E. H., Dager, S., Estes, A., Gallagher, L., Malow, B. A., Parr, J. R., Spence, S. J., Vorstman, J., Frey, B. J., Robinson, J. T., Strug, L. J., Fernandez, B. A., Elsabbagh, M., Carter, M. T., Hallmayer, J., Knoppers, B. M., Anagnostou, E., Szatmari, P., Ring, R. H., Glazer, D., Pletcher, M. T. & Scherer, S. W. 2017. Whole genome sequencing resource identifies 18 new candidate genes for autism spectrum disorder. Nat Neurosci, 20, 602-611.

Yuen, R. K., Merico, D., Cao, H., Pellecchia, G., Alipanahi, B., Thiruvahindrapuram, B., Tong, X., Sun, Y., Cao, D., Zhang, T., Wu, X., Jin, X., Zhou, Z., Liu, X., Nalpathamkalam, T., Walker, S., Howe, J. L., Wang, Z., Macdonald, J. R., Chan, A., D’abate, L., Deneault, E., Siu, M. T., Tammimies, K., Uddin, M., Zarrei, M., Wang, M., Li, Y., Wang, J., Wang, J., Yang, H., Bookman, M., Bingham, J., Gross, S. S., Loy, D., Pletcher, M., Marshall, C. R., Anagnostou, E., Zwaigenbaum, L., Weksberg, R., Fernandez, B. A., Roberts, W., Szatmari, P., Glazer, D., Frey, B. J., Ring, R. H., Xu, X. & Scherer, S. W. 2016. Genome-wide characteristics of de novo mutations in autism. NPJ Genom Med, 1, 160271-1602710.

Yuen, R. K., Thiruvahindrapuram, B., Merico, D., Walker, S., Tammimies, K., Hoang, N., Chrysler, C., Nalpathamkalam, T., Pellecchia, G., Liu, Y., Gazzellone, M. J., D’Abate, L., Deneault, E., Howe, J. L., Liu, R. S., Thompson, A., Zarrei, M., Uddin, M., Marshall, C. R., Ring, R. H., Zwaigenbaum, L., Ray, P. N., Weksberg, R., Carter, M. T., Fernandez, B. A., Roberts, W., Szatmari, P. & Scherer, S. W. 2015. Whole-genome sequencing of quartet families with autism spectrum disorder. Nat Med, 21, 185-91.

Zhang, J., Kinch, L. N., Cong, Q., Weile, J., Sun, S., Cote, A. G., Roth, F. P. & Grishin, N. V. 2017. Assessing predictions of fitness effects of missense mutations in SUMO-conjugating enzyme UBE2I. Hum Mutat, 38, 1051-1063.

Zhang, X., Tamaru, H., Khan, S. I., Horton, J. R., Keefe, L. J., Selker, E. U. & Cheng, X. 2002. Structure of the Neurospora SET domain protein DIM-5, a histone H3 lysine methyltransferase. Cell, 111, 117-27.

Zhang, X., Yang, Z., Khan, S. I., Horton, J. R., Tamaru, H., Selker, E. U. & Cheng, X. 2003. Structural basis for the product specificity of histone lysine methyltransferases. Mol Cell, 12, 177-85.

Zhang, Y., Pak, C., Han, Y., Ahlenius, H., Zhang, Z., Chanda, S., Marro, S., Patzke, C., Acuna, C., Covy, J., Xu, W., Yang, N., Danko, T., Chen, L., Wernig, M. & Sudhof, T. C. 2013. Rapid single-step induction of functional neurons from human pluripotent stem cells. Neuron, 78, 785-98.

Zylicz, J. J., Dietmann, S., Gunesdogan, U., Hackett, J. A., Cougot, D., Lee, C. & Surani, M. A. 2015. Chromatin dynamics and the role of G9a in gene regulation and enhancer silencing during early mouse development. Elife, 4.

